# Multimodal learning of noncoding variant effects using genome sequence and chromatin structure

**DOI:** 10.1101/2022.12.20.521331

**Authors:** Wuwei Tan, Yang Shen

## Abstract

**Motivation:** A growing amount of noncoding genetic variants, including single-nucleotide polymorphisms (SNPs), are found to be associated with complex human traits and diseases. Their mechanistic interpretation is relatively limited and can use the help from computational prediction of their effects on epigenetic profiles. However, current models often focus on local, 1D genome sequence determinants and disregard global, 3D chromatin structure that critically affects epigenetic events.

**Results:** We find that noncoding variants of unexpected high similarity in epigenetic profiles, with regards to their relatively low similarity in local sequences, can be largely attributed to their proximity in chromatin structure. Accordingly we have developed a multimodal deep learning scheme that incorporates both data of 1D genome sequence and 3D chromatin structure for predicting noncoding variant effects. Specifically, we have integrated convolutional and recurrent neural networks for sequence embedding and graph neural networks for structure embedding despite the resolution gap between the two types of data, while utilizing recent DNA language models. Numerical results show that our models outperform competing sequence-only models in predicting epigenetic profiles and their use of long-range interactions complement sequence-only models in extracting regulatory motifs. They prove to be excellent predictors for noncoding variant effects in gene expression and pathogenicity, whether in unsupervised “zero-shot” learning or supervised “few-shot” learning.

**Availability:** Codes and data access can be found at https://github.com/Shen-Lab/ncVarPred-1D3D

**Supplementary information:** Supplementary data are available at *Bioinformatics* online.

## 1 Introduction

Genetic variation in the noncoding regions (estimated to constitute 98–99% of human genomes) can be linked to traits and diseases (van Ouwerkerk *et al*., 2019; Lim and Kim, 2019; Ohnmacht *et al*., 2020; Frydas *et al*., 2022). Growing data and analyses from genome-wide association studies (GWAS) help identify and interpret such links (Zhang and Lupski, 2015), along with evolving computational methods. GWAVA (Ritchie *et al*., 2014) prioritized noncoding variants by machine learning from various functional annotations (largely by the ENCODE Project Consortium). FunSeq2 (Fu *et al*., 2014) did so for cancer somatic variants using a weighted scoring scheme. Then epigenetic impact prediction was introduced for mechanistic interpretations, whereas deep learning models emerged. To predict DNA accessibility (specifically, DNase I hypersensitive sites), Basset (Kelley *et al*., 2016) applied convolutional neural networks (CNN) to learn from the data annotated by ENCODE (ENCODE Project Consortium, 2012) and Roadmap Epigenomics (Roadmap Epigenomics Consortium *et al*., 2015). To predict transcription factor binding sites and histone marks besides DNase I hypersensitive sites, DeepSEA (Zhou and Troyanskaya, 2015) also used CNN. Focused on transcription factor binding, DanQ (Quang and Xie, 2016) combined CNN and recurrent neural networks (RNN; specifically, a bidirectional long short-term memory or bidirectional LSTM) to capture the long term dependencies between motifs, and TBiNet (Park *et al*., 2020) further included an attention mechanism for model interpretability. Recently a transformer, developed for long sequences using a sparse self-attention mechanism, was also applied to the topic (Zaheer *et al*., 2020).

As surveyed above, current machine learning methods for epigenetic impacts of noncoding variants only treat DNA as 1D sequences (often local ones up to 1,000 base pairs or 1K bps). Although correctly focused on sequence determinants (Koohy *et al*., 2013; Boeva, 2016; Ngo *et al*., 2019), they disregard that DNA sequences are packed in 3D chromosome structure whereas such 3D global perspective of DNA sequences is important to regulatory mechanisms and functional annotation of noncoding variants (Klemm *et al*., 2019; D’haene and Vergult, 2021). Technologies such as Hi-C (Lieberman-Aiden *et al*., 2009) and ChIA-PET (Fullwood *et al*., 2009) reveal chromatin interactions such as long-range intra- and inter-chromosome enhancer–promoter interactions (Zhang *et al*., 2013; Dai *et al*., 2018; Chen *et al*., 2020) controlling gene expression.

In this study we fill the gap by incorporating both 1D local sequences and 3D global structures into predicting epigenetic impacts of noncoding variants. The approach of multimodal data integration is currently lacking for the topic, although some recent methods predicted how noncoding variant sequences can change chromatin conformations (Trieu *et al*., 2020; Zhou, 2022). Specifically, we first statistically test the hypothesis underlying the approach and verify that disparity between the levels of sequence (feature) similarity and epigenetic profile (label) similarity can be attributed to chromatin interactions. Given the rationale verified we then introduce a series of models combining the embedding of 1D local sequences (through CNN, RNN, and transformer) and the embedding of 3D global structures / interaction intensity graphs (through various graph neural networks, or GNN), while overcoming the challenge from modality difference (texts versus graphs) and resolution difference (1-bp versus 100K-bp). We show that our models outperform DeepSEA and DanQ in predicting epigenetic profiles of genome sequences on held-out chromosomes. And we analyze the (in)sensitivity of the improvements with regards to the cell line and the resolution associated with the input Hi-C data. We further examine our models’ interpretability in extracting regulatory motifs and found that long-range interactions in chromatin structure data enhanced such interpretability. We last demonstrate utility of our models for predicting noncoding variant effects in gene expression (eQTL) and pathogenicity, using model-predicted epigenetic profile changes as zero-shot or few-shot predictors of variant effects.

## 2 Materials and methods

### 2.1 Data

#### DNA sequences and epigenetic profiles

We used the same data set processed as in DeepSEA (Zhou and Troyanskaya, 2015). The data set contains 2.6 million non-overlapping 200-bp bins on either strand (approximately 17% of the human genome) that is described by a 1K-bp neighborhood encompassing the 200-bp bin at its center and is doubled considering both strands of DNA. Each of the resulting 5.2 million samples is labelled with binary values for 919 epigenetic events associated with 148 cell lines, including binding to 690 transcription factors from 91 cell lines, 125 DNase I hypersensitive sites from 119 cell lines, and 104 histone marks from 8 cell lines. We also used the same data split as in DeepSEA, that is, chromosome 8 and 9 (445 thousand samples on both strands, i.e., 227,512 bins on either) for testing, part of chromosome 7 (8,000 samples) for validation, and the rest (4.4 million samples) for training.

#### Chromatin structures

We additionally used Hi-C data from the ENCODE portal (Luo *et al*., 2020), in the form of interaction frequency matrices, that map genome-wide chromatin interactions. Cell lines GM12878, IMR90 and K562 were related to the 919 epigenetic events. More details can be found in see Supplementary Sec. S1.

### 2.2 Hypothesis Tests on the Significance of 3D Information

For a given cell line’s given replicate, we randomly drew 0.2% of the samples; partitioned sample pairs with non-zero frequencies into three sets whose sequence similarity percentiles were over, around, or under profile similarity profiles, respectively; and performed one-side Kolmogorov– Smirnov (K-S) test to compare the interaction frequencies of the “under” (or “over”) set and those of the “consistent” set (the null hypothesis being that the former are below (or above) the latter). The process was repeated 100 times and the frequencies of p-value falling below 0.05 were collected. More details can be found in Supplementary Sec. S2.

### 2.3 Machine Learning for Predicting Epigenetic Profiles

#### Architecture

As shown in the schematic illustration in Figure 1, our models take two inputs, a 1D local sequence of one kilobase and a 2D global interaction-frequency matrix of the whole genome. The two types of inputs are separately encoded with multi-layer neural networks, except in our most advanced versions (to be detailed next), to address a resolution gap — whereas local DNA sequences are available in every base, global chromosome interactions are typically among regions of 100 kilobases. The resulting embeddings are concatenated and fed to a fully-connected layer with 919 neurons and sigmoid activation functions that outputs the probability of each of the 919 epigenetic events.

**Fig. 1:**
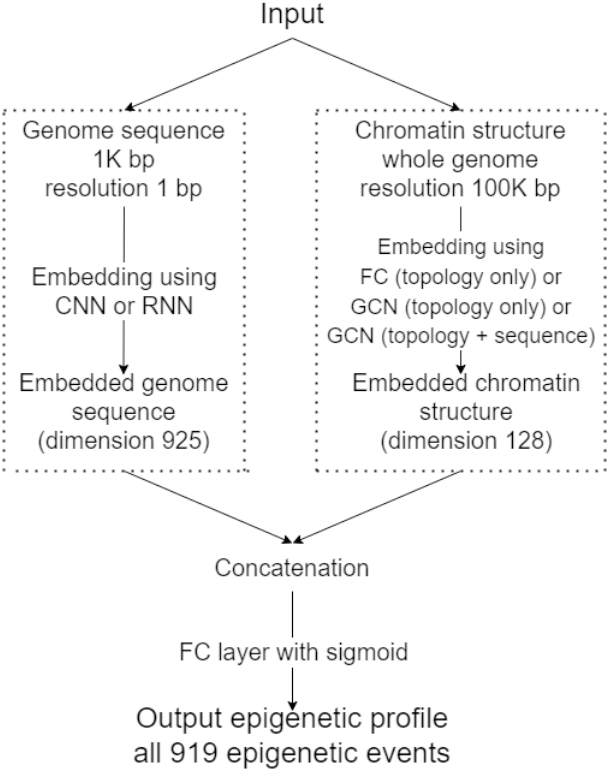
The illustration of our proposed models for epigenetics that can be used for zero-shot or few-shot prediction of noncoding variant effects.

For local sequence embedding (left branch in Figure 1) of each nucleotide, we used two versions: 3 layers of 1D CNNs and 2 layers of max pooling interspersed as in DeepSEA as well as 1 layer of CNN and 1 layer of max pooling followed by 1 layer of biLSTM as in DanQ. Whereas 1D CNNs are good at extracting and combining local sequence features, biLSTM can capture long-term dependencies. Using a 1D CNN before biLSTM also helps reduce the input size and allieviate the resulting training difficulties for biLSTM. We intentionally kept their architectures the same as DeepSEA and DanQ, respectively, to isolate the contribution of chromatin structures. Resulting sequence embeddings were of 925 dimensions for each kilobase.

For global structure embedding (right branch in Figure 1) for each chromatin region, we used three versions where the first two consider chromatin topologies only and the last additionally considers DNA sequences. (1) MLP (topology only): Depending on the chromatin region (100 kilobase long for Hi-C of a 100K-bp resolution) that it belonged to, each kilobase sample was featurized by the corresponding row / column of the chromatin interaction frequency matrix (after normalization) and encoded through a multi-layer perception (MLP), specifically 3 fully connected (FC) layers of 1000, 400, and 128 neurons, respectively, each followed by ReLU activation. (2) GCN (topology only): Each kilobase sample was treated as a node in a graph whose node features were simply all ones of 768 dimensions and edge weights were corresponding normalized interaction frequencies. It was encoded through graph convolutional networks or GCN (Kipf and Welling, 2016) of 3 layers (latent dimensions being 1000, 400, and 128, respectively). Whereas MLPs can capture global relationship across the chromatin, GNNs including GCNs better capture graph structures and their sparsity. We note that most of the Hi-C data used in the study have less than 10% nonzero frequencies. (3) GCN (sequence+topology): We additionally introduced DNA sequences to the GCN (topology only) encoder by replacing the all-one node features with mean-pooled regional sequence embedding from a pretrained DNA language model, DNABERT (Ji *et al*., 2021). Specifically, we used DNABERT to embed each 100K-bp with a 500-bp sliding window and a 250-bp stride by averaging the 768-dimensional embeddings of all windows. For each window, we used 497 tokens (495 6-mers plus special tokens CLS for sentence class and SEP as sentence separator), obtained a 768 by 497 dimensional embedding, and used the row average to mitigate the impact of the starting position.

More details about model architectures are in Supplementary Sec. S3.1

#### Training

We used binary cross entropy with L1 and L2 regularization as the training loss and optimized hyperparameters including learning rate, the two regularization coefficients, and the choice of replicate per cell line (that provides chromatin structure to encode) over combinations of options. For each hyperparameter combination, part of the models were warm-started with DNA sequence encoders (CNN or CNN/RNN) initialized at the values from pretrained sequence-only models (DeepSEA or DanQ) and trained for up to 40 epochs with early-stopping (patience being 4 epochs). The optimal hyperparameters were chosen based on validation loss. In particular, replicates ENCFF014VMM, ENCFF928NJV, and ENCFF013TGD were chosen for cell lines GM12878, IMR90, and K562, respectively, in CNN+MLP and fixed in our other models. Once hyperparameters were tuned, each model was trained 5 times with the cold-started portion randomly initialized and its performances’ means and standard deviations were based on the 5 repeats. More details about model training including values of tuned hyperparameters can be found in the Supplementary Sec. S3.2.

### 2.4 Interpreting the Learned Epigenetic Predictors

Following DanQ (Quang and Xie, 2016), we examined the regulatory significance of the 320 kernels in the first convolutional layer for sequence embedding (kernel size being 8 and 26 bases for CNN and CNN/RNN, respectively). Specifically, for each kernel, the contiguous 8 or 26 bases with the largest activation value were selected for each test sequences and a position-specific frequency matrix (profile) was accordingly built over all test sequences. We queried each of the 320 model-learned profiles against HOCOMOCO (v11 core) (Kulakovskiy *et al*., 2018), a dataset of human transcription-factor binding motifs, using the tool TOMTOM (Bailey *et al*., 2015) with a cutoff E < 0.01.

### 2.5 Noncoding Variant Effect Prediction: eQTL

We use the trained epigenetic predictors as featurizers to predict noncoding variant effect on expression levels, that is, to predict eQTL (expression quantitative trait loci). We collected cell line specific eQTL data from the Genotype-Tissue Expression project (GTEx Analysis v7): 3845 entries for GM12878 and 11091 for IMR90 using the filter q-value no greater than 0.05, labeled with whether gene expression levels increase or decrease. These two sets of data were relatively label balanced – positive (increase) rates were 49.6% for GM12878 and 49.8% for IMR90. For each entry, trained epigenetic predictors (sequence-based DeepSEA and DanQ as well as our sequence and cell line specific structure-based models described above) calculated the differences in predicted probabilities (between the wild type and variants) and the differences in predicted log odds for cell-line epigenetic events (91 for GM12878 and 5 for IMR90 among 919). Using the predicted differences as features, we held out 10% data for hyper-parameter tuning and used the rest for 10-fold cross validation when building logistic regression models with l2 regularization (scikit-learn). Through 10-fold cross validation AUROC over the 10% held out data, we chose the differences in log odds as features and tuned the values for the inverse of regularization strength (Supplementary Sec. S7). As each epigenetic predictor (such as CNN+MLP) was trained five times and used as fixed encoders, the corresponding supervised eQTL classifier was accordingly trained five times to calculate the average AUROC and AUPRC over 10 folds of the 90%. In the end, we report the mean and the standard deviation of the average performances for each model.

### 2.6 Noncoding Variant Effect Prediction: Pathogenecity

We used ncVarDB (Biggs *et al*., 2020), a manually curated noncoding variant data set, to assess our models’ performance in identifying pathogenic variants. Out of 721 pathogenic variants and 7226 benign variants in ncVarDB, 713 pathogenic and 7134 benign variants were retained as the test set (positive rate: 9.08%), 20 variants as the validation set, and 80 variants as the source of the training set that grows from 0 to 80 at the increment of 10, simulating zero-shot (unsupervised) to few-shot (supervised) learning. In the case of zero-shot unsupervised learning where no ncVarDB data is used, we directly used the absolute difference in the predicted epigenetic probabilities (finally chosen for the better performance) or the difference in log odds, between the wild-type/reference and the alternate/variant sequences, as a pathogenicity indicator. In the case of few-shot supervised learning, we used the pre-trained epigenetic predictors to build a Siamese neural network: two identical genome sequence-structure encoders, one for wild-type / reference and one for variant / alternative sequences, whose squared differences of the 919 epigenetic probabilities, magnitude-only and differentiable, are fed to a 919D-to-1D FC layer followed by a sigmoid activation function, to predict whether the given variant is pathogenic or benign. The resulting pathogenicity classifier was trained end to end (unlike the eQTL classifier where a single encoder is used and fixed), using 10, 20, …, 80 training samples, with all parameters warm-started but the last FC layer. The loss function was balanced cross entropy with L2 regularization whose weights were tuned from 1E-12 to 1E-8 log-uniformly and optimized at 1E-11 with validation loss. As each type of epigenetic predictor (such as CNN+MLP) was trained five times, the corresponding pathogenicity classifier was accordingly initialized and trained five times before an average performance is reported.

## 3 Results

In this section, we first statistically assess the hypothesis that the inconsistency between genome sequence similarity and epigenetic profile similarity can be partly attributed to the chromatin structure. With the hypothesis verified, we adopted it as the rationale of our approach and started with the simplest version of models where chromatin structures are encoded with fully connected layers. We showed that this straightforward way to introduce chromatin structures already outperformed the competing sequence-only models; and confirmed that the model improved the performances by correcting the bias from genome sequence alone using chromatin structure. We also examined the (in)sensitivity of the model to the resolution and the cell line of the input chromatin structure data.

We further evaluate the performances of the more advanced versions where chromatin interactions are modeled as graphs and region sequences as nodes are embedded by DNA language models. These models are more accurate in epigenetic profile prediction than competing sequence-only models; and they are explainable in extracting regulatory motifs. Importantly, as demonstrated in eQTL and pathogenicity prediction, these models are well-performing unsupervised zero-shot predictors of noncoding variant effects; and they can be flexibly fine-tuned to broadly applicable and accurate few-shot predictors, using few labeled data and no feature engineering.

### 3.1 Chromatin structure helps explain a gap between DNA sequence similarity and epigenetic profile similarity

Current machine-learning methods predict epigenetics based on DNA sequences alone, thus leading to similar epigenetics predictions for similar DNA sequences. However DNA sequence similarity and epigenetic profile similarity are not always consistent and sometimes in disparity. To explain the disparity, we hypothesize that sample pairs with low (high) DNA sequence similarity but high (low) epigenetic profile similarity tend to be closer (farther) in 3D, compared to those with consistent sequence and epigenetic similarities. Two-sample one-sided K-S tests validated the hypothesis (Figure 2) in all but few bio replicates (Hi-C experiment data for chromatin structure). The outliers with no statistical significance included three bio replicates of cell line GM12878, two of which (replicates 6 and 12) were also outliers with the most dense interactions (possibly including more spurious weak interactions). More details can be found in the Supplementary Section S2.

**Fig. 2:**
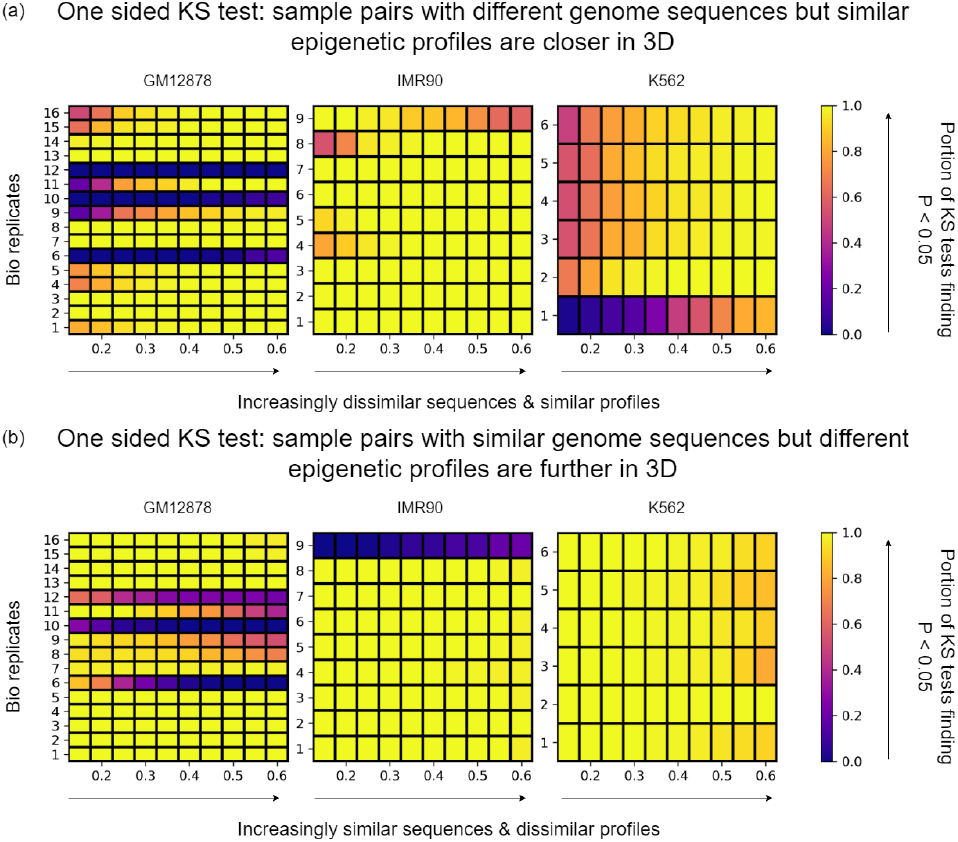
Sequence pairs whose sequence similarity percentiles (a) exceed or (b) lag epigenetic-profile similarity percentiles tend to be closer or further in 3D chromatin structure (across bio-replicates or Hi-C data of chromatin structures for cell lines GM12878, IMR90, and K562), which is frequently found with statistical significance (p-value below 0.05). The trend intensifies when sequence pairs are increasingly dissimilar in sequence but similar in epigenetic profiles. Few outliers exist and some can be attributed to denser but possibly spurious interactions in those replicates.

For sample pairs with low sequence high profile similarity, the trend (as measured by the portions of trials with statistical significance) was increasing as the sequence–profile disparity (as measured by the cumulative percentage difference or margin between the two similarities) increased. In contrast, sample pairs with high sequence low profile similarity did not see more significance as the disparity increased. The observations agree with biological knowledge: spatial proximity helps explain high profile similarity unexpected from low sequence similarity, as it implies similar chromosome access for epigenetic factors in transcription or modification; but spatial remoteness doesn’t necessarily explain the opposite disparity, as it doesn’t necessarily determine dissimilar chromosome access.

### 3.2 Our models outperform sequence-only epigenetic predictors by filling the gap with chromatin structure

The statistical significance of chromatin structures in addressing the sequence–profile disparity suggests that the additional input of chromatin structure besides DNA sequences would improve epigenetic prediction. We used the chromatin structure data from each of the three cell lines (GM12878, IMR90, and K562) to predict the whole epigenetic profile from 148 cell lines. And we combined two encoders (CNN and CNN/RNN) for local DNA sequences and three encoders (MLP, GCN [topology only] and GCN with DNABERT [sequence and topology]). Due to label imbalance (positive rate merely 2.06%), we focused on AUPRC (area under the precision-recall curve) as the major assessment metric and included AUROC (area under the receiver operating characteristic curve) in the Supplementary Table S2.

All our models outperformed DeepSEA and DanQ (Table 1). The most advanced version using GCN with DNABERT to encode chromatin sequences and structures improved AUPRC by over 10% compared to DeepSEA or reproduced DanQ. We note that the most advanced CNN/RNN+GCN with DNABERT encodes the chromatin’s global sequence and structure together and is complemented by local sequence embedding for desired sensitivity within chromatin regions. The overall improvements were not very sensitive to the input structures’ cell lines especially when GCNs were used. They did slightly decreased (within 0.01) if the resolution of the chromatin structure or the dimension of its embedding decreased (Supplementary Sec. S5.1 and S5.2). They were relatively insensitive to parameter initialization and normalization during model training (Supplementary Sec. S5.3).

**Table 1.**
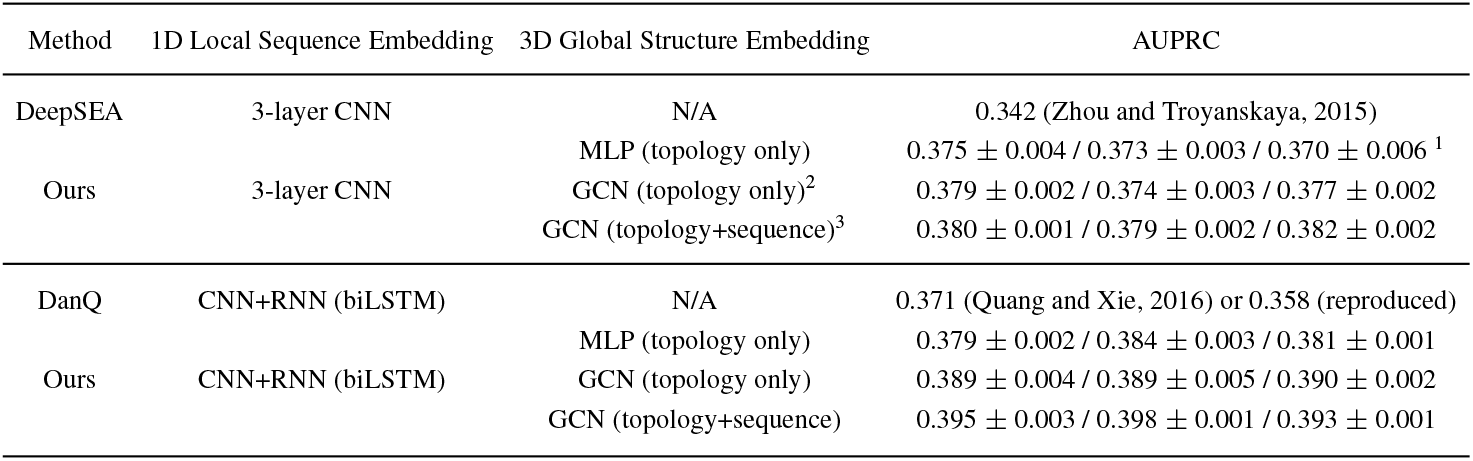
Epigenetic profile prediction assessed in AUPRC (Area Under the Precision-Recall Curve) whose base-line value for random classifiers is 0.02. Our models intentionally used the same neural network architectures for DNA local 1D sequence embedding as in DeepSEA or DanQ. Thus their additional introduction of chromatin global 3D structure embedding led to much improved and robust performances. ^1^ Performances using chromatin structure data from the cell line GM12878 / IMR90 / K562, respectively. ^2^ All-one node features for 100K-bp regions. ^3^ DNABERT-encoded node features for 100K-bp regions.

We next examine the origin of the improvements. To remove possible distraction of model architectures and isolate the contribution of chromatin structures, we compared our basic model, CNN+MLP, to DeepSEA (based on the same CNN for local sequence embedding). We calculated the Spearman correlations between the true and the predicted label similarities for sample pairs randomly picked in the test set (as opposed to random pairs from all the data for hypothesis tests in Sec. 3.1) and reported the distribution of the Spearman correlations over aforementioned 100 trials in Figure 3. The chromatin structure information improved the correlation in all three subsets, whether sequence similarity levels were under, consistent with, or over profile similarity levels. And it did the most (+0.05) for DNA pairs whose sequence similarities were significantly under profile similarities, by correcting the sequence-only bias with structure information. Statistical significance of the improvements were supported by one-sided K-S tests (Supplementary Sec. S4.1 including Tables S2–S4).

**Fig. 3:**
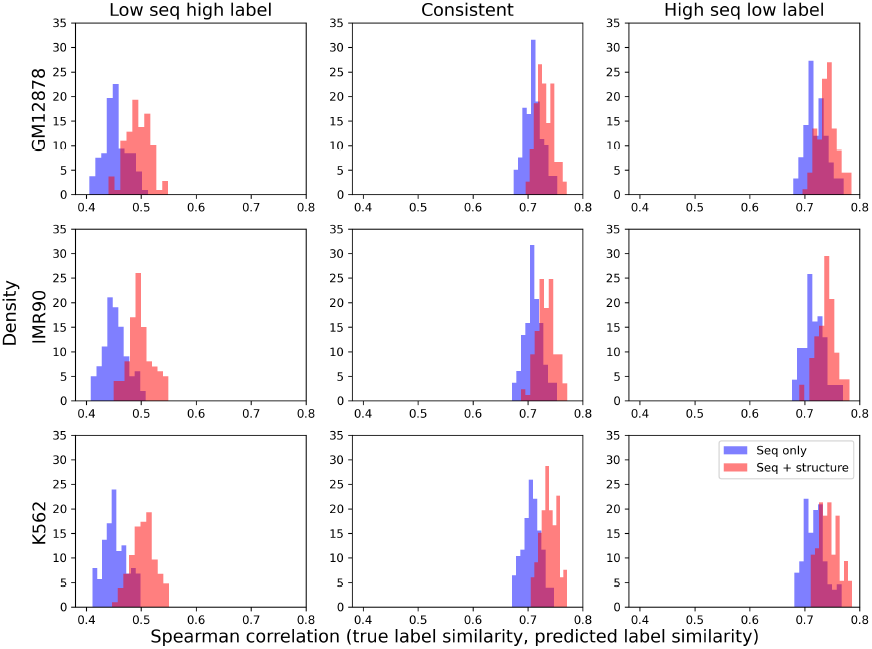
Comparing sequence-only DeepSEA and our sequence+structure basic model, CNN+MLP, in how much they address the disparity between sequence similarity and epigenetic profile similarity. For random sample pairs in each subset / column of a disparity level, we calculate the Spearman correlation between actual and predicted epigenetic profile similarities; and we repeat the process 100 times and compare the Spearman correlation distributions between DeepSEA and our CNN+MLP while using the chromatin structure of each cell line / row.

### 3.3 Our models benefit from cell line specificity but tolerate otherwise

After verifying and tracing the help of chromatin structure (cell line dependent) for epigenetic prediction, we examine whether more help is there for epigenetic events matching the chromatin structure’s cell line or specific types of epigenetic events.

We evaluated AUPRC (AUROC) values for 919 individual epigenetic events’ predictions and reported the percentages of the events where CNN+MLP outperformed DeepSEA (Table 2 and Supplementary Figure S3). Overall, 89%–94% of all events saw more accurate predictions with the additional input of any chromatin structure (among the three). When the statistics were broken down based on cell line match, matching epigenetic events had around 6–9% more frequencies than other events to enjoy such accuracy boost. Therefore, our epigenetic predictors, being cell line specific, benefit from such cell line specificity.

**Table 2.** Percentages of the 919 epigenetic events whose CNN+MLP predictions outperformed DeepSEA (CNN) in AUROC or AUPRC.

| Hi-C cell line | % with better AUROC |  |  | % with better AUPRC |  |  |
| --- | --- | --- | --- | --- | --- | --- |
|  | GM12878 | IMR90 | K562 | GM12878 | IMR90 | K562 |
| Same cell line events | 98.02 | 100 | 96.79 | 99.12 | 96.00 | 96.05 |
| Other events | 92.33 | 93.84 | 87.80 | 92.29 | 94.15 | 87.70 |
| All events | 92.90 | 93.88 | 89.39 | 92.96 | 94.16 | 89.17 |

We further examined the AUPRC boost from DeepSEA to our CNN+MLP for the epigenetic events of non-matching cell lines and checked whether the extent of the boost correlated with the degree of cell line similarity. Specifically, we used the CNN+MLP trained with chromatin structure of K562 (ENCFFF013TGD) and checked the AUPRC boost for some of the 919 epigenetic events associated with 8 more cell lines. Their identities, expression cosine similarity (based on TPM [transcripts per million] values from Dependency Map ^1^), and the numbers of associated epigenetic events can be found in the texts in Figure 4 along with violin plots of AUPRC boosts (due to the use of chromatin structure for K562). The violin plots were based on five times the number of associated epigenetic events’ performance differences, because our models were trained in 5 trials. A weak Spearman correlation of 0.28 was found between median AUPRC boosts and expression similarity to cell line K562, albeit without statistical significance (p-value 0.46).

**Fig. 4:**
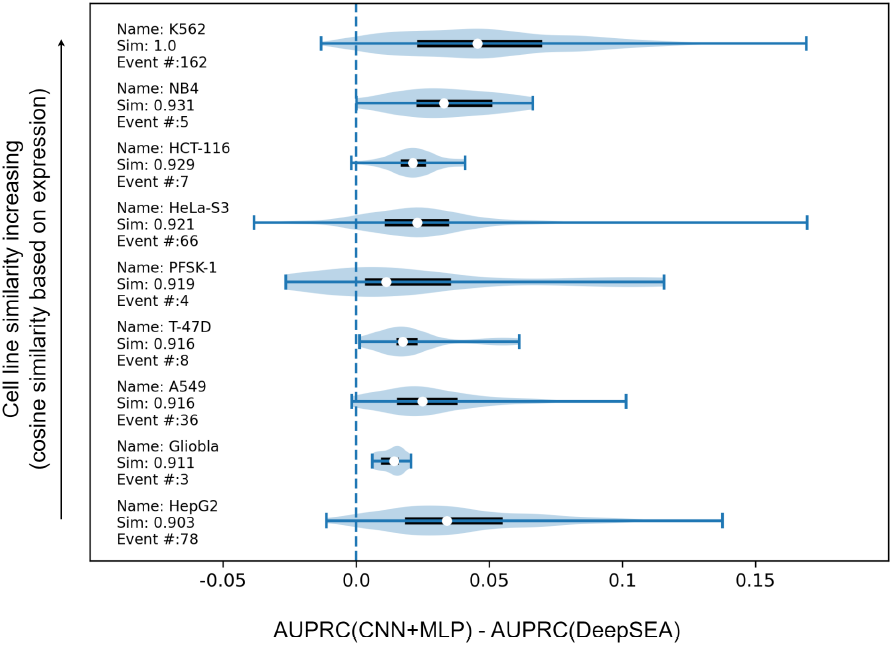
Violin plots of performance boost from DeepSEA to our CNN+MLP (trained with K562 chromatin structure) for epigenetic events of various cell lines. The white dot, the box, and the whiskers indicate the median, the 25%–75% percentile, and the extremes, respectively. The cell lines are arranged in the order of expression similarity to the input K562 cell line, with names, cosine similarities, and numbers of associated epigenetic events given below corresponding violin plots.

We also checked the sensitivity of the trained model to the chromatin structure usage. All available chromatin structures, 52 Hi-C experiments from 10 cell lines are collected. We used the pretrained fixed models and made inference on the test set with all available Hi-C. Figure S6 in the Supplemental Sec. S5.4 shows that mismatched cell line’s chromatin structure input didn’t negatively impact the performance for an average epigenetic event, and it did for those events related to the cell line whose chromatin structure was used during training. Nevertheless, either way our basic CNN+MLP performed better than sequence-only DeepSEA when tested with a mismatched cell line’s chromatin structure input.

### 3.4 Our models are interpretable in capturing regulatory motifs

Following DanQ, we analyzed key regions that our models identified over test sequences and compared them to known sequence motifs for transcription factor (TF) binding. Out of 320 convolutional kernels of size 26, 164–167 kernels (depending on the cell line of the input chromatin structure) in our basic CNN/RNN+MLP were found to match at least one known sequence motifs, which led to 166–168 motifs (see details in the Supplemental Sec. S6.1). Similar numbers were reported in our CNN/RNN+GCN without or with DNABERT. The numbers of matched kernels and identified motifs in our models were similar to those in DanQ (169 matched kernels and 172 motifs).

Our chromatin structure-informed models showed complementarity to sequence-only DanQ. When examining motifs identified by CNN/RNN+GCN without DNABERT (thus global chromatin embedding only used topology and not DNA sequences), we found 2 motifs that were missed by DanQ but consistently identified by our model, regardless of which cell line’s chromatin structure was used. These two motifs are related to genes POU3F1 (both strands) and TFDP1 (Figure 5). The discovery of these two motifs was attributed to the use of chromatin structure that suggests long-range interactions. More details can be found in the Supplemental Sec. S6.3. Comparing our CNN/RNN+MLP to DanQ led to similar observations.

**Fig. 5:**
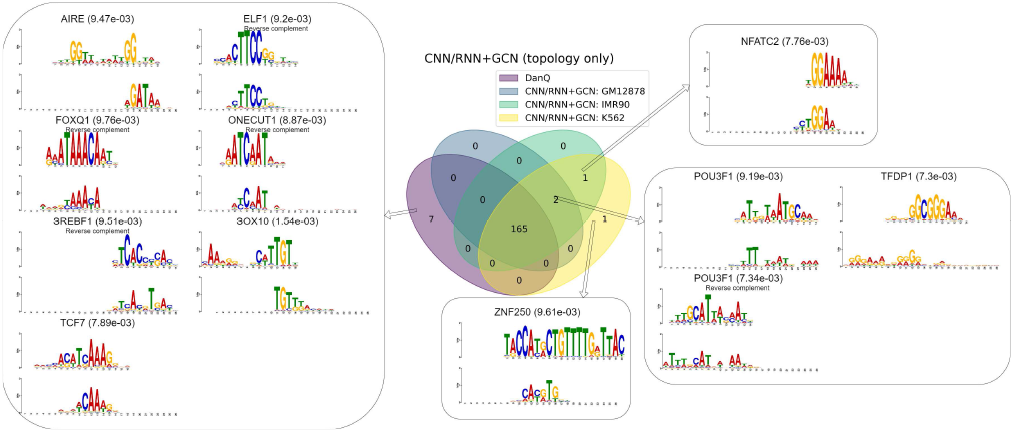
Our CNN/RNN+GCN model identified 2 motifs (lower right) that were missed by DanQ, regardless of which of the three Hi-C inputs were used. They missed 7 motifs (left) that lacked long-range interactions and had low interaction frequencies.

Meanwhile, there were 7 motifs found by sequence-based DanQ but missed by all three of our models using CNN/RNN to embed local DNA sequences, no matter which neural networks was used to embed global chromatin structure (and sequence). Whereas we believe that high-resolution chromatin structure data might mitigate the issue, we found that these 7 motifs lacked long-range interactions mechanistically and were of lower interaction frequencies compared to other motifs (Supplementary Sec. S6.4)

### 3.5 Noncoding Variant Effect Prediction: eQTL

We used the aforementioned models trained to predict epigenetic events as fixed, feature generators and trained supervised machine learning models to predict noncoding variant effect on gene expression (increase or decrease).

For cell line specific, noncoding eQTL data from GTEx (3845 samples for GM12878 and 11091 for IMR90), cell line specific features (91 for GM12878 and 5 for IMR90) were used to train regularized logistic regression models. Figure 6 shows that, compared to using epigenetic predictions from sequence-only DeepSEA, using those from our basic model to utilize chromatin structure, CNN+MLP, was on average (over 5 trials) 0.19*σ* worse (in AUROC) / 0.62*σ* better (in AUPRC) for GM12878 and 2.83*σ* better / 3.77*σ* better for IMR90 (*σ* is the standard deviation of our model’s performances over five training trials), when focusing on the eQTLs that caused significant expression changes (fold change cutoff at 2^2^). The margins over DeepSEA-predicted epigenetic features improved when the focus shift to eQTLs with even more significant expression changes (fold change cutoff at 2^2.5^): 0.58*σ* / 1.37*σ* better for for GM12878 and 2.52*σ* / 3.69*σ* better for IMR90. We similarly compared epigenetic features predicted by DeepSEA to those by our CNN+GCN with DNABERT and found that similar conclusions held and CNN+GCN with DNABERT (where chromatin structure and sequence were jointly embedded) further improved the margins (Supplementary Figure S13). We also compared epigentic predictions by DanQ to those by our CNN/RNN+GCN without or with DNABERT (Supplementary Figure S13 and S14) and reached similar conclusions. Therefore, the input of chromatin structure improved the prediction of epigenetic events, which further improved eQTL prediction.

**Fig. 6:**
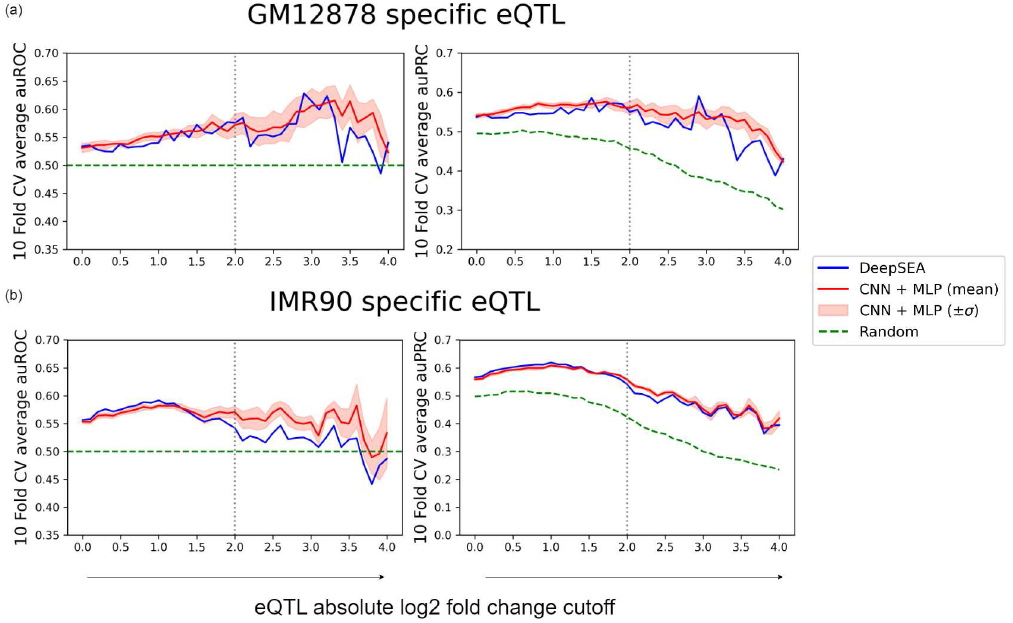
eQTL prediction performances of DNA sequence-only DeepSEA and our chromatin structure-informed CNN+MLP (mean and standard deviation over 5 trials), at various cutoff of actual expression level fold changes for the test set. Note that as the cutoff of expression change further increases, the number of eQTLs involved further decreases until being too low to support statistical significance. So our analysis ended at the fold change cutoff of 2^4^.

**Fig. 7:**
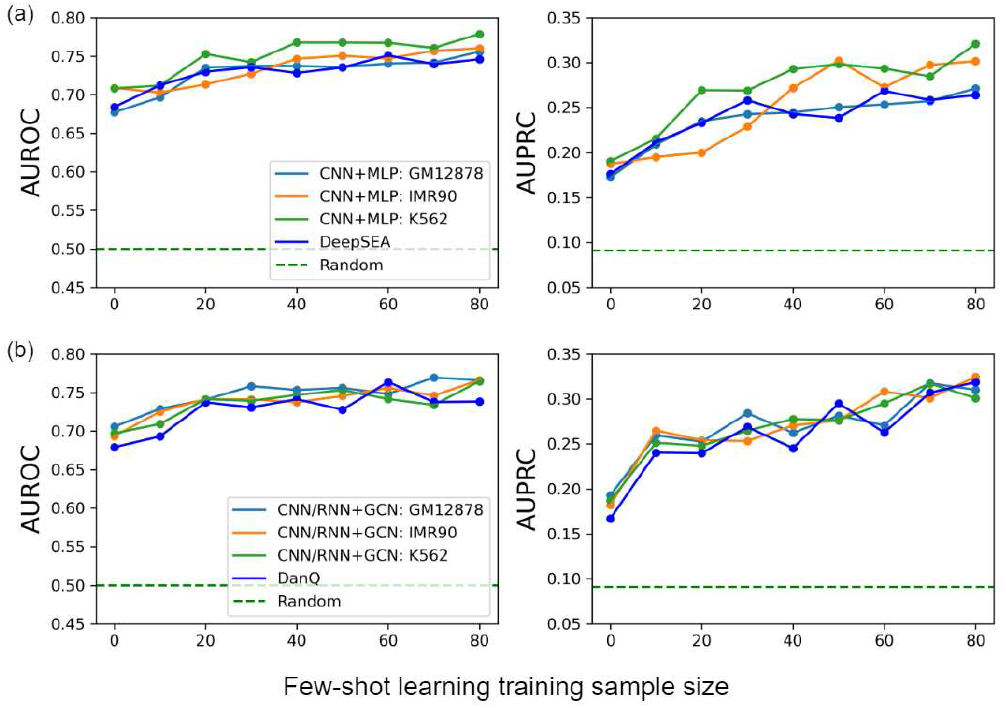
Test performances (AUROC and AUPRC) of zero-shot or few-shot pathogenicity predictors, using (a) CNN+MLP and (b) CNN/RNN+GCN (with DNABERT), improve as the size of the training set increases.

### 3.6 Noncoding Variant Effect Prediction: Pathogenicity

We also tried to assess our models in identifying pathogenic versus benign noncoding variants, whether directly used as an unsupervised “zero-shot” learner or incorporated into a Siamese neural network to be a supervised “few-shot” learner. Figure shows that, even without access to labeled data (variants with known pathogenicity or benign), our zero-shot pathogenicity predictors was of AUROC around 0.7 and AUPRC around 0.18. With as few as tens variants, our few-shot pathogenicity predictors quickly improved the performances as the training data grow and reached AUROC over 0.75 and AUPRC over 0.3. Such improvements were consistent across model versions or Hi-C data used. The more advanced CNN/RNN+GCN (with DNABERT) showed even more consistency across Hi-C data choices. Numerical values on the unsupervised performances are in the Supplementary Sec. S8.1.

Our noncoding variant effect predictors are broadly applicable to substitutions, insertions, and deletions across various genome positions. In fact, state of theart predictors CADD (Rentzsch *et al*., 2021), DANN (Quang *et al*., 2015) and FATHMM-XF (Rogers *et al*., 2018), failed to make prediction for 535 of the 7847 test variants, including those on chromosomes X, Y, and M as well as some substitutions across various genome positions, which makes it impossible to compare performances fairly. This was in contrast to our models as detailed in the Supplementary Sec. S8.2.

## 4 Conclusion

Toward noncoding variant effect prediction, we have built multi-modal machine learning models that use both genome sequences and chromatin structures to predict epigenetic profiles, whereas previous methods only used genome sequences. To rationalize the approach, we have verified that there is often a disparity between sequence similarity and epigenetic-profile similarity and such disparity can be explained by, thus mitigated by, spatial proximity. Accordingly, we have used the combination of various neural networks to embed local 1D kilobase sequences (CNN and CNN/RNN) and global 3D chromosome structures represented in 2D graphs (MLP and GCN), that are available in different resolutions, and concatenated the embeddings to feed epigenetic prediction layers. We have additionally introduced a version where the two embedding processes are coupled, by using a pretrained DNA language model to embed 100K-long bases as node features before embedding the 2D graphs of chromatin structures. Compared to sequence-only methods, our models robustly improve the accuracy of epigenetic prediction and complement the explainability in extracting regulatory motifs.

For noncoding variant effect prediction, these pre-trained models are used directly as unsupervised zero-shot predictors or incorporated into a Siamese neural network as supervised few-shot predictors, as demonstrated in eQTL and pathogenicity prediction. Without manual feature engineering, our models are shown to be fine-tunable for various effects and broadly applicable to variants of various types at various genome positions.

## Acknowledgement

This project was in part supported by the National Institute of General Medical Sciences (R35GM124952 to Y.S.). Portions of this research were conducted with the advanced computing resources provided by Texas A&M High Performance Research Computing.

## Footnotes

1 https://doi.org/10.6084/m9.figshare.19139906.v1

