## Supplementary Data for "Multimodal learning of noncoding variant effects using genome sequence and chromatin structure"

Wuwei Tan and Yang Shen
Texas A&M University

December 2022

#### 1 Hi-C Experiment Data Summary

We downloaded Hi-C data in August 2022 from the ENCODE portal [1] using the following filters of experiment search — organism: 'Homo sapiens', perturbation: 'not perturbed', genome assembly: 'hg19' (as in DeepSEA's data), and available file types: 'hic'. Out of ten resulting cell lines, three were related to the 919 epigenetic events, namely GM12878, IMR90 and K562. With these cell lines chosen for the biosample filter, we downloaded corresponding Hi-C data with quality status as 'released' and output type as 'mapping quality threshold chromatin interactions', including 16, 9, and 6 bio replicates (experiments) for GM12878, IMR90 and K562, respectively. We used the 100K-bp resolution by default and also obtained that of 500K and 1M bps for analyses.

There were 52 Hi-C experiments from 10 cell lines as of the end of August 2022 as follows:

- Endothelial cell of umbilical vein: ENCFF606XNW
- **GM12878**: ENCFF014VMM, ENCFF053BXY, ENCFF065LSP, ENCFF223UBX, ENCFF227XJZ, ENCFF355OWW, ENCFF473CAA, ENCFF482LGO, ENCFF514XWQ, ENCFF563XES, ENCFF632MFV, ENCFF688KOY, ENCFF718AWL, ENCFF777KBU, ENCFF799QGA, ENCFF812THZ
- GM23248: ENCFF768UBD
- HAP1: ENCFF230HVV
- hTERT RPE-1: ENCFF922ERE
- **IMR90**: ENCFF029MPB, ENCFF043EEE, ENCFF303PCK, ENCFF366ERB, ENCFF894GLR, ENCFF920CJR, ENCFF928NJV, ENCFF997RGL, ENCFF999YXX
- KBM-7: ENCFF239BHZ, ENCFF277LAN, ENCFF397CMD, ENCFF698KFV, ENCFF945TUH
- Keratinocyte: ENCFF349RZY, ENCFF406KJN, ENCFF569RJM, ENCFF738YON
- **K562**: ENCFF013TGD, ENCFF097SKJ, ENCFF406HHC, ENCFF464KRA, ENCFF929RPW, ENCFF996XEO

- Mammary epithelial cell: ENCFF198SSL, ENCFF251UEF, ENCFF307PDL, ENCFF491AOR, ENCFF543USQ, ENCFF706SFK, ENCFF773ITV, ENCFF942LTN

Bold-faced are three cell lines associated with some of the 919 epigenetic events to predict, and the order of the replicates for each cell line corresponds to their indices in the subsequent hypothesis tests, such as **Figure 2** of the main text.

The Hi-C data used are in the form of interaction frequency matrices. Higher interaction frequencies in the 2D data indicate closer proximity in 3D. The raw interaction frequency matrices were normalized by adding one's to its diagonal elements and then dividing each element by the square root of the product of the row sum and the column sum [2].

### 2 Hypothesis Tests on the Significance of 3D Information

#### 2.1 Subsets of sequence pairs defined by sequence–profile disparity

For a given pair of genome sequences $\mathbf{x}_i$ and $\mathbf{x}_j$ (of lengths $L = 1000$ in our study), the sequence similarity is defined as $\text{SIM}_{\text{seq}}(\mathbf{x}_i, \mathbf{x}_j) = \frac{1}{L}\sum_{l=1}^{L} \mathbb{1}\{x_i^l = x_j^l\}$ where $x_i^l$ ($x_j^l$) is the categorical type of the $l$-th nucleic acid in the sequence $\mathbf{x}_i$ ($\mathbf{x}_j$). For a given pair of epigenetic profiles (labels) $\mathbf{y}_i = \mathbf{y}(\mathbf{x}_i)$ and $\mathbf{y}_j = \mathbf{y}(\mathbf{x}_j)$ (of 919 dimensions in our study) for sequences $\mathbf{x}_i$ and $\mathbf{x}_j$, the epigenetic profile similarity is defined as $\text{SIM}_{\text{epigen}} = \frac{1}{919}\sum_{k=1}^{919} \mathbb{1}\{y_i^k = y_j^k\}$ where $y_i^k$ ($y_j^k$) is the binary value for the $k$-th epigenetic event in the profile $\mathbf{y}_i$ ($\mathbf{y}_j$).

Within a selected set of sequence pairs (among 0.2% random samples of sequences we chose all pairs with non-zero normalized interaction frequencies), we calculated the cumulative percentage (percentile rank) for the sequence similarity and that for the profile similarity, for each pair $\mathbf{x}_i$ and $\mathbf{x}_j$: $\text{CumuPct}(\text{SIM}_{\text{seq}}(\mathbf{x}_i, \mathbf{x}_j))$ and $\text{CumuPct}(\text{SIM}_{\text{epigen}}(\mathbf{x}_i, \mathbf{x}_j))$. For instance, if a given pair has similarity above 30% of all pairs considered, then its cumulative percentage is 30. Accordingly, among all pairs considered, the subsets where sequence similarity is under, around, and over profile similarity are defined as:
$\{(\mathbf{x}_i, \mathbf{x}_j) | \text{CumuPct}(\text{SIM}_{\text{seq}}(\mathbf{x}_i, \mathbf{x}_j)) \leqslant \text{CumuPct}(\text{SIM}_{\text{epigen}}(\mathbf{x}_i, \mathbf{x}_j)) - \delta\}$,
$\{(\mathbf{x}_i, \mathbf{x}_j) | \text{CumuPct}(\text{SIM}_{\text{epigen}}(\mathbf{x}_i, \mathbf{x}_j)) - \delta \leqslant \text{CumuPct}(\text{SIM}_{\text{seq}}(\mathbf{x}_i, \mathbf{x}_j)) \leqslant \text{CumuPct}(\text{SIM}_{\text{epigen}}(\mathbf{x}_i, \mathbf{x}_j)) + \delta\}$, and
$\{(\mathbf{x}_i, \mathbf{x}_j) | \text{CumuPct}(\text{SIM}_{\text{seq}}(\mathbf{x}_i, \mathbf{x}_j)) \geqslant \text{CumuPct}(\text{SIM}_{\text{epigen}}(\mathbf{x}_i, \mathbf{x}_j)) + \delta\}$, respectively.
$\delta$ is a margin parameter. The "under" and "over" subsets were also referred to as low-sequence high-profile similarity and high-sequence low-profile similarity, respectively.

#### 2.2 Data visualization

We illustrate the three subsets for the interaction frequency data, using the example of Hi-C experiment ENCFF014VMM for cell line GM12878, the same replicate chosen my our machine learning models for the cell line (see details in the next section). In Figure S1 each pair is shown as a point in the 2D space of cumulative percentages in sequence and profile similarities, and colored with the cumulative percentages in normalized interaction frequencies. Visual inspection indicated that the "low-sequence high-profile similarity" ("under") region was enriched with higher interaction frequencies (closer proximity).

#### 2.3 Kolmogorov-Smirnov tests

To rigorously compare those pairs with low-sequence high-profile similarity (or high-sequence low-profile similarity) and those pairs with consistent similarities, 100 statistical tests (one-sided two-

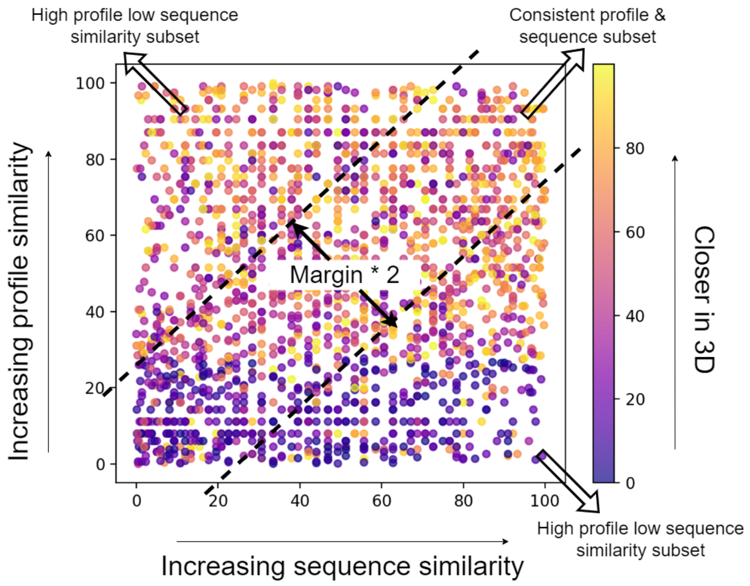


Figure S1: Three subsets of sequence pairs, defined by the disparity between their percentiles in sequence similarity ($x$-axis) and percentiles in epigenetic profile similarity ($y$-axis), show distinct patterns in their 3D proximity / normalized interaction frequencies (colored in cumulative percentage). Sequence pairs whose percentiles in epigenetic profile similarity significantly surpass their percentiles in sequence similarity (upper left corner) tend to be enriched with closer proximity (brighter color).

sample Kolmogorov-Smirnov tests) were employed, following 100 trials of initial 0.2% samples, for each Hi-C data corresponding to a replicate (experiment) of a given cell line. The null hypotheses were that the normalized interaction frequencies of the low-sequence high-profile similarity (high-sequence low-profile similarity) subset were no higher (lower) than those of the consistent sequence-profile similarity subset. We evaluated how often p-values were below 0.05 among the 100 trials.

From Figure 2 of the main text we found that all but few replicates showed statistical significance of dominant frequencies especially when the separating margins increased, supporting that those pairs with exceptionally low (high) epigenetic profile similarities compared to their sequence similarities tended to be further (closer) in 3D.

#### 2.4 Outliers

We traced some origins of the few outliers in the K-S tests above. First, we examined the sparsity of the normalized interaction frequencies for all involved Hi-C replicates by defining sparsity as the portion of the zero elements in a normalized interaction frequency matrix. We found that the three Hi-C replicate outliers for cell line GM12878 were of lower sparsity compared to most other replicates for the same cell line, including two with the exceptionally lowest sparsity (Figure S2 left). However this was not the case for cell line IMR90. For sequence pairs sampled in all GM12878 replicates we further calculated the Spearman correlation between the sparsity and the K-S statistic (the distance between empirical distribution functions), as a function of sequence–profile disparity margin $\delta$. We found that the correlations were positive and high for both "under" and "over" subsets (Figure S2 right), supporting that low sparsity for this cell line contributes to low statistical significance. The "over" subset (high sequence low profile/label similarity) seemed to be more sensitive to sparsity, judging by its higher Spearman correlations. Meanwhile, the Spearman correlation for the "under" subset increased as $\delta$ increased, showing that the standing out interactions dominated the sparsity–disparity relationship for these sequences. So if the low sparsity was partially due to spurious weak interactions misidentified in experiments, increasing margin tended to spot the standing-out, strong interactions more. In contrast, the Spearman correlation for the "over" subset was relatively stable as $\delta$ changed.

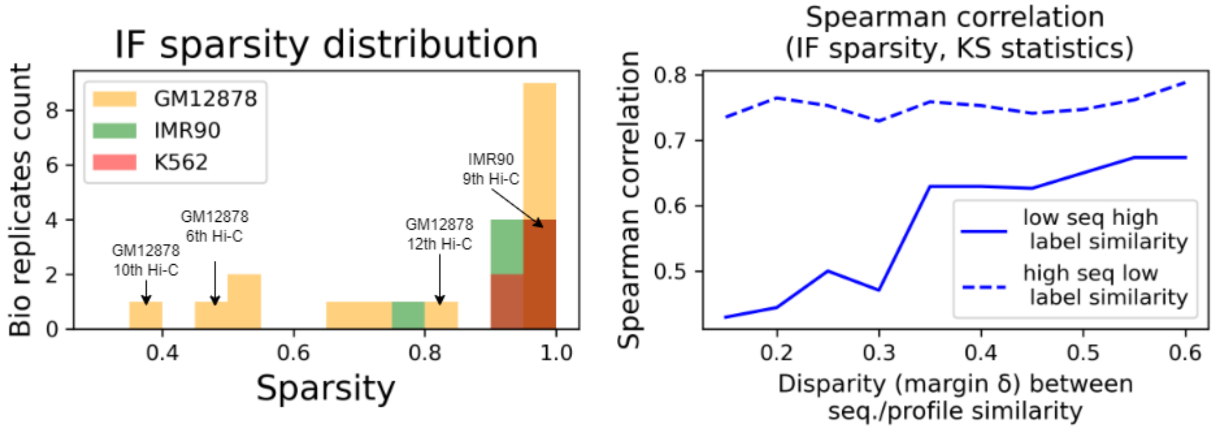


Figure S2: (Left) The histograms of Hi-C experiment data's 3D interaction frequency sparsity levels. Arrow pointed are few outliers found in K-S tests and they tend to be of low sparsity for cell line GM12878. (Right) For sequence pairs sampled for cell line GM12878, the Spearman correlation between 3D interaction sparsity and K-S statistic was positive and high for both "under" (low sequence high label similarity) and "over" (high sequence low label similarity) and it increased for the "under" subset as the sequence–profile disparity margin $\delta$ increased. The results suggest that, as epigenetic profile similarity exceedingly surpasses sequence similarity, Hi-C data of less non-zero interaction frequencies possibly due to reduced spurious weak interactions would better reveal such disparity.

### 3 Machine Learning for Epigenetic Profile Prediction

#### 3.1 Model architectures

**Local DNA sequence embedding.** To encode local DNA sequences (kilobases) we intentionally used the model architectures from state-of-the-art sequence-only epigenetic predictors as follows.

CNN as in DeepSEA [3]:

1. Convolution layer (320 kernels, kernel size: 8, step size 1) with ReLU
2. Max pooling layer (kernel size 4, step size 4)
3. Dropout (20 percent)
4. Convolution layer (480 kernels, kernel size 8, step size 1) with ReLU
5. Max pooling layer (kernel size 4, step size 4)
6. Dropout (20 percent)

7. Convolution layer (960 kernels, kernel size 8, step size 1) with ReLU
8. Dropout (50 percent)
9. Fully connected layer (925 neurons)

CNN/RNN as in DanQ [4]:

1. Convolutional layer (320 kernels, kernel size 26, step size 1) with ReLU
2. Pooling layer (Window size 13, step size 13)
3. Dropout (20 percent)
4. BiLSTM layer (containing 320 hidden features)
5. Dropout (50 percent)
6. Fully connected layer (925 neurons)

**Global chromatin structure embedding.** We additionally used the following three architectures to embed chromatin regions (100 kilobases) where each kilobase belongs to.

MLP (topology only):

1. Fully connected layer (1000 neurons) with ReLU. The input dimension is varies with the chromatin structure resolution – 30971 for 100K bp, 6207 for 500K bp and 3114 for 1M bp.
2. Fully connected layer (400 neurons) with ReLU
3. Fully connected layer (128 neurons) with ReLU

GCN (topology only):

1. GCN layer (input node feature [all-ones] dimension 768, output node feature dimension 1000) with ReLU.
2. Dropout (20 percent)
3. GCN layer (output node feature dimension 400)
4. Dropout (20 percent)
5. GCN layer (output node feature dimension 128)

GCN (sequence + topology):

1. GCN layer (input node feature [mean-pooled DNABERT over 497 tokens first and then over all 500-bp windows] dimension 768, output node feature dimension 1000) with ReLU.
2. Dropout (20 percent)
3. GCN layer (output node feature dimension 400)
4. Dropout (20 percent)
5. GCN layer (output node feature dimension 128)

**Epigenetic profile prediction (Output).** Based on aforementioned local sequence and global structure embedding, we predict the probability of each event in the epigenetic profile, using the following architecture.

1. Concatenation layer to combine the local sequence embedding of a kilobase (925 dimensions) and the global chromatin structure embedding of the 100 kilobases that it belong to (128 dimensions) followed by ReLU
2. Fully connected layer (919 neurons) with sigmoid

#### 3.2 Model training

The loss function includes binary cross entropy as well as L1 (LASSO) and L2 (Ridge) regularization. Binary cross entropy was averaged over all training samples in a given batch, positive (sense) or negative (antisense) strand, that were essentially treated independent. L1 regularization was on parameters of the chromatin structure encoder (right branch of Figure 1 in the main text) and those of the output fully connected layer; and L2 regularization was on all parameters of each model.

We used the Adam optimizer with default weight decay schedule to train our models, using data in batches of size 512, for up to 40 epochs unless early stopping criteria is met (see below). We initialized the DNA-sequence encoders not randomly but from pre-trained DeepSEA and DanQ for CNN and CNN/RNN respectively. The pre-trained DeepSEA was directly downloaded from the original publication's shared data; and the pre-trained DanQ was reproduced in PyTorch by ourselves (the original release was in Keras). The other parameters were initialized randomly and the impact of these parameters' initialization was studied in Sec. S5.3. Early stopping criteria is a patience of 4 epochs (40 validations, see below) based on the validation loss. In other words, before reaching the maximum epochs, training would stop if no decrease is observed for validation loss in four consecutive epochs. The checkpoint with the lowest validation loss was chosen to define optimal parameters of our models.

Our hyperparameters include the initial learning rate for the optimizer Adam, the coefficients of L1 and L2 regularizations, and the bio replicate (defining which input to feed the chromatin structure encoder) for a given cell line. For each input Hi-C replicate option for the three cell lines described in Sec. S1, we performed grid search over these hyperparameters ($lr \in \{$1E-2, 5E-3, 1E-3, 5E-4, 1E-4, 5E-5, 1E-5$\}$, L1 coefficient $\in \{$1E-6, 1E-7, ..., 1E-12$\}$, L2 coefficient $\in \{$1E-6, 1E-7, ..., 1E-12$\}$) and trained model parameters for each hyperparameter combination. We chose the optimal hyperparameters based on the validation loss averaged over all input Hi-C replicates. We calculated validation loss 10 times equi-spaced during each epoch, each time using the incremental 10% training data to train and all validation data to validate. The tuned hyperparameters can be found in Table S1. With the tuned hyperparameters, we chose the optimal replicates for each cell line: in the CNN+MLP model they were ENCFF014VMM for cell line GM12878, ENCFF928NJV for cell line IMR90, and ENCFF013TGD for cell line K562; and the same replicates were used for our other models.

With hyperameters tuned, each of our models was trained five times with random initialization (see more details below) and used to make inference on the test set. Following DeepSEA, our trained models predicted probabilities of epigenetic events for each pair of samples corresponding to the same 1K-bp on two strands and took the average as the final prediction for the 1K-bp. AUPRC and AUROC were calculated over all testing 1K-bp's and their mean and standard deviation over 5 training repeats were reported for each model.

Table S1: Tuned hyperparameters including learning rate and regularization weights.

| | Learning rate | L1 weight | L2 weight |
|---|---|---|---|
| CNN+MLP | 5E-5 | 1E-11 | 1E-8 |
| CNN+GCN w/ all-ones | 5E-5 | 1E-12 | 1E-8 |
| CNN+GCN w/ DNABERT | 5E-5 | 1E-12 | 1E-8 |
| CNN+GCN w/ DNABERT (binary graph) | 1E-4 | 1E-11 | 1E-9 |
| CNN/RNN + MLP | 5E-5 | 1E-11 | 1E-9 |
| CNN/RNN+GCN (topology only) | 5E-5 | 1E-12 | 1E-8 |
| CNN/RNN+GCN w/ DNABERT | 1E-4 | 1E-12 | 1E-8 |
| CNN+MLP (resolution 500K bp) | 5E-5 | 1E-11 | 1E-10 |
| CNN+MLP (resolution 1M bp) | 5E-5 | 1E-12 | 1E-10 |
| CNN+MLP (embedding 64) | 5E-5 | 1E-12 | 1E-10 |
| CNN+MLP (embedding 256) | 5E-5 | 1E-11 | 1E-8 |
| CNN+MLP (embedding 512) | 5E-5 | 1E-11 | 1E-10 |

##### 3.3 Performance comparison

Besides using the major assessment metric of AUPRC (Area Under the Precision-Recall Curve) in Table 1 of the main text, we also used another metric of AUROC (Area Under the Receiver Operating Characteristic curve) to compare our models' performances in epigenetic profile prediction with the state of the art (DeepSEA and DanQ). We intentionally used the same neural network architectures for DNA local 1D sequence embedding as those in DeepSEA and DanQ, so that performance margins can be attributed surely to the newly added chromatin global 3D structure embedding. Due to the extreme class imbalance (positive rate merely 2.06%), AUPRC in Table 1 is a more appropriate assessment metric and AUROC below tended to be saturated.

Table S2: Epigenetic profile prediction assessed in AUROC (Area Under the Receiver Operating Characteristic curve) whose base-line value for random classifiers is 0.50. Our models intentionally used the same neural network architectures for DNA local 1D sequence embedding as in DeepSEA or DanQ and their additional introduction of chromatin global 3D structure embedding led to improved and robust performances. [1]Performances using chromatin structure data from the cell line GM12878 / IMR90 / K562, respectively. [2]All-one node features for 100K-bp regions. [3]DNABERT-encoded node features for 100K-bp regions.

| Method | 1D Local Sequence Embedding | 3D Global Structure Embedding | AUROC |
|---|---|---|---|
| DeepSEA | 3-layer CNN | N/A | 0.933 [3] |
| | | MLP (topology only) | $0.941 \pm 0.001$ / $0.940 \pm 0.000$ / $0.939 \pm 0.002$ |
| Ours | 3-layer CNN | GCN (topology only)[2] | $0.942 \pm 0.001$ / $0.940 \pm 0.001$ / $0.942 \pm 0.001$ |
| | | GCN (topology + sequence) [3] | $0.942 \pm 0.000$ / $0.942 \pm 0.000$ / $0.943 \pm 0.001$ |
| DanQ | CNN + RNN (biLSTM) | N/A | 0.938 [4] / 0.930 (reproduced) |
| | | MLP (topology only) | $0.938 \pm 0.000$ / $0.939 \pm 0.001$ / $0.939 \pm 0.001$ |
| Ours | CNN + RNN (biLSTM) | GCN (topology only) | $0.940 \pm 0.001$ / $0.940 \pm 0.001$ / $0.941 \pm 0.000$ |
| | | GCN (topology + sequence) | $0.941 \pm 0.001$ / $0.942 \pm 0.000$ / $0.941 \pm 0.000$ |

### 4 Tracing the origin of improvements by comparing DeepSEA (sequence-only) and our CNN+MLP (sequence + structure)

In an effort to trace the origin of our models' improvements and verify the contribution of our models' rationale (tested in Section S2), we examined which subset of test samples or sample pairs benefited more compared to others. We compared DeepSEA and our basic CNN+MLP for this purpose.

#### 4.1 Which subset of sample pairs with sequence–profile disparity

Similar to our hypothesis tests in Section S2, we randomly drew 1% of the test samples, chose all pairs but those with zero interaction frequencies, and calculated their sequence similarity, (actual) label/profile similarity, and predicted label similarity (1 less the difference in predicted probability, averaged over all 919 epigenetic events / labels). We used sequence–profile disparity margin $\delta = 20\%$ (so separation is $2\delta = 40\%$, see Fig. S1) to partition sampled pairs into three roughly equal-sized subsets. And we compared performances of DeepSEA and our CNN+MLP in the three subsets, using one-sided K-S tests and Jensen-Shannon distances. Tables S3, S4, and S5 show that improvements were made among all 3 cell lines and all 3 subsets (low sequence high label similarity or "under", consistent sequence and label similarity, and high sequence low label similarity or "over"). The largest improvement was for the "under" subset, indicating that long range interactions in chromatin 3D structure data are picked up in our models to boost prediction similarity for sequentially far yet structurally close DNA sequences.

Table S3: CNN+MLP using GM12878 chromatin structure versus DeepSEA

| | low seq high label | consistent | high seq low label |
|---|---|---|---|
| one-sided KS test statistics | 6.38E-1±4.53E-2 | 4.96E-1±6.41E-2 | 4.22E-1±5.27E-2 |
| one-sided KS test p value | 2.50E-16±4.89E-16 | 1.56E-8±3.11E-8 | 1.69E-6±3.18E-6 |
| JS distance | 8.81E-3±6.18E-4 | 3.15E-3±2.06E-4 | 4.30E-3±6.18E-5 |

Table S4: CNN+MLP using IMR90 chromatin structure versus DeepSEA

| | low seq high label | consistent | high seq low label |
|---|---|---|---|
| one-sided KS test statistics | 7.48E-1±5.23E-2 | 6.32E-1±9.72E-2 | 5.12E-1±8.03E-2 |
| one-sided KS test p value | 1.33E-22±2.67E-22 | 8.45E-12±1.69E-11 | 8.00E-8±1.60E-7 |
| JS distance | 8.77E-3±2.63E-4 | 3.35E-3±2.39E-4 | 4.63E-3±3.64E-4 |

Table S5: CNN+MLP using K562 chromatin structure versus DeepSEA

| | low seq high label | consistent | high seq low label |
|---|---|---|---|
| one-sided KS test statistics | 7.14E-1±3.00E-2 | 5.78E-1±5.78E-2 | 4.44E-1±5.24E-2 |
| one-sided KS test p value | 1.34E-22±2.66E-22 | 1.00E-12±2.00E-12 | 3.77E-7±7.50E-7 |
| JS distance | 9.73E-3±7.61e-4 | 3.58E-3±1.27E-4 | 4.75E-3±2.07E-4 |

#### 4.2 Which subset of epigenetic events: cell lines and event types

We did head-to-head comparison between DeepSEA and our CNN+MLP in AUROC or AUPRC and showed the scatter plots in Figure S3 where each symbol corresponded to one of the 919 epigenetic events to predict. We differentiated the symbols based on cell lines and event types and asked which subset of epigenetic events benefited more thanks to the introduction of input chromatin structure data. Results indicated that epigenetic events of the same cell line as that of the input chromatin structure and epigenetic events of transcription factor binding were among the biggest beneficiaries.

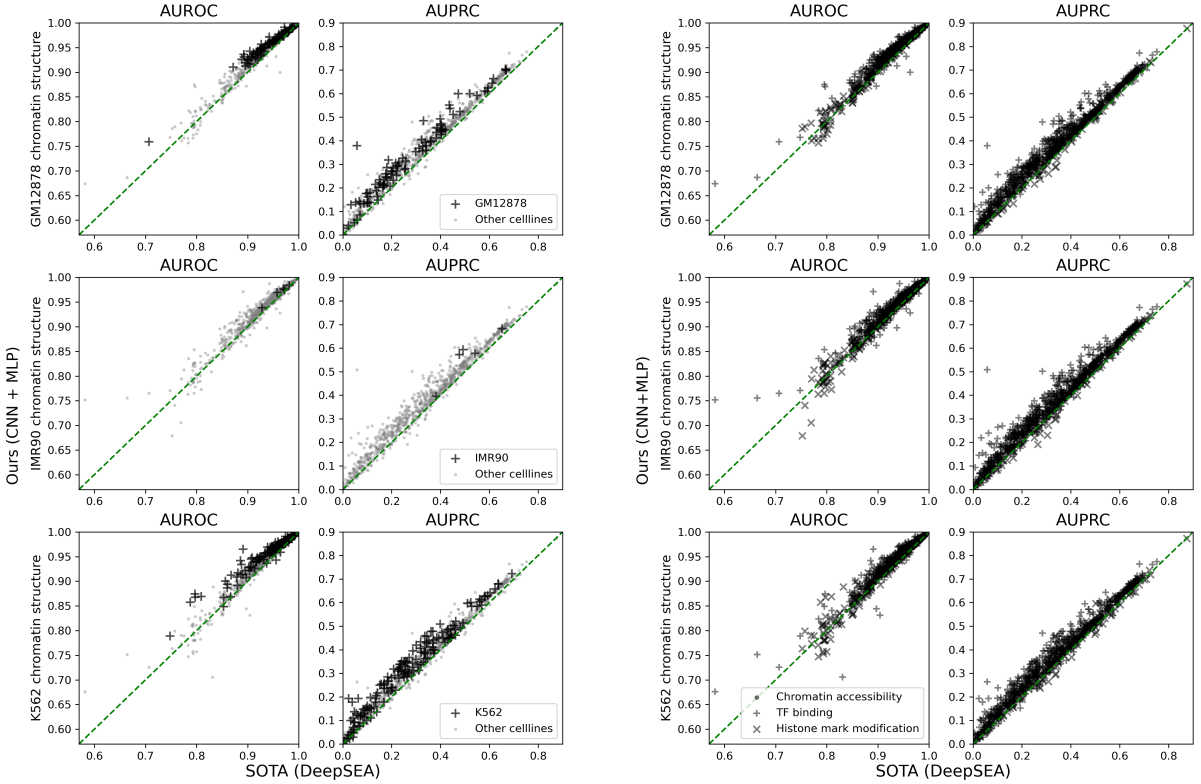


Figure S3: (Left) Using CNN+MLP to incorporate additional input of chromatin structure, epigentic profile prediction improved more against DeepSEA for epigenetic events associated with the same cell line as that of the input chromatin structure. (Right) The most extreme improvements were often for epigenetic events related to transcription factor binding.

### 5 Sensitivity Analysis of Model Performances

We also tested on how sensitive our models are to input data, model architecture, and model training, using CNN+MLP as an example.

#### 5.1 Impact of model input: resolution of chromatin structure Hi-C data

To analyze the impact of the input chromatin structure's resolution, we used same cell lines' data of lower resolutions (500K-bp and 1M-bp) to train the model and only changed the dimension of the input layer for chromatin structure embedding accordingly. As shown in Figure S4, model performances were relatively stable with regard to input chromatin structure's resolution and consistently above DeepSEA. Higher resolution of 100K-bp did perform better and even higher resolutions could help even more.

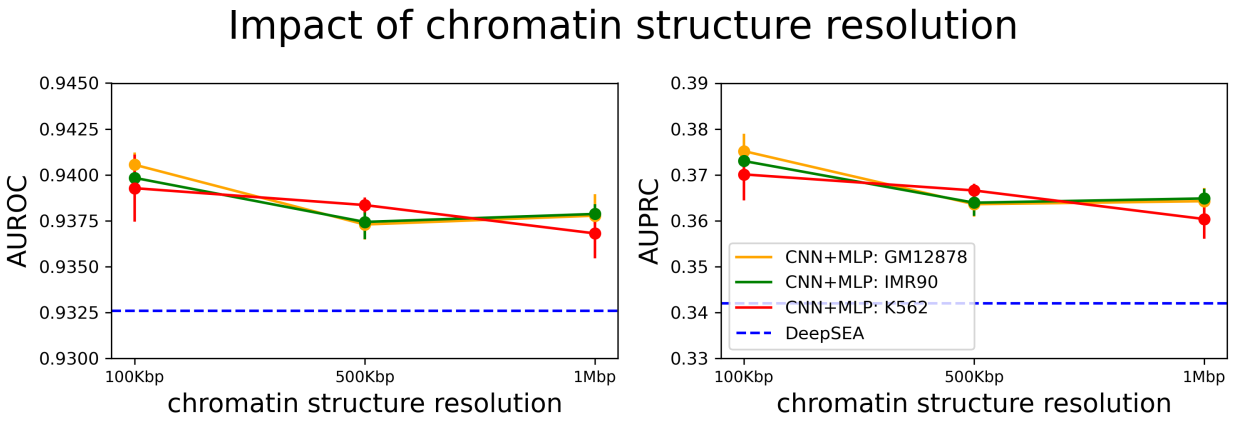


Figure S4: Epigenetic prediction performances of our CNN+MLP while using the input of different cell lines' chromatin structures of different resolutions. They all outperformed sequence-only DeepSEA (dashed line) and higher resolution (100K-bp) helped.

##### 5.2 Impact of model architecture: dimension of chromatin embedding

With input chromatin structure's resolution fixed at 100K-bp, we tested on the dimension of the latent space embedding the input structure. We varied the output dimension of the last fully connected layer of MLP for chromatin embedding to 64, 128 (default), 256, and 512 and kept using the previously determined optimal hyperparameters. Figure S5 shows relative stable performances consistently outperforming DeepSEA, regardless of the embedding dimension. Reducing the latent dimension of chromatin structure embedding from the default 128 to 64 noticeably lowered the performances whereas increasing it to 256 or 512 did not help either. The results show a tricky balance between sequence embedding and structure embedding especially when DNA sequences and chromatin structures are of different resolutions.

##### 5.3 Impact of model training: parameter initialization and normalization

We have tested the impact of changing the parameter initialization or adopting batch normalization, without changing the model architecture. In our default setting, the parameters of the fully connected layers in the chromatin embedding and after the concatenation of embeddings were initialized following the uniform distribution $U(-\frac{1}{\sqrt{k}}, \frac{1}{\sqrt{k}})$, where k is the number of the features. Here we have additonally tested the Xavier uniform initialization with the gain parameter (for scaling) at 1 as well as the Xavier normal initialization with gain at 1.0, which are implemented in PyTorch. We also tested adding batch normalization for all fully connected layers for chromatin structure embedding (right branch in Figure 1 of the main text). We used the same L1 and L2 regularization weights and the same learning rate, as the optimal values in the default CNN+MLP, for the tests.

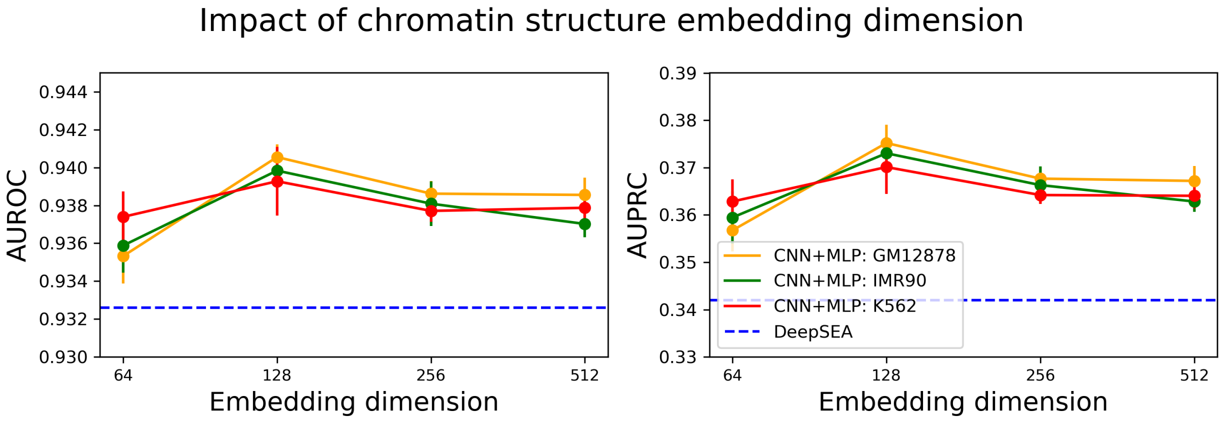


Figure S5: Epigenetic prediction performances of our CNN+MLP while using the input of different cell lines' chromatin structures embedded in different dimensions. They all outperformed sequence-only DeepSEA (dashed line).

Table S6: Our CNN+MLP models' AUROC using different initializers or using batch normalization.

| | GM12878 | IMR90 | K562 |
|---|---|---|---|
| default initialization | 0.9406±0.0007 | **0.9398 ± 0.0004** | 0.9393±0.0018 |
| xavier uniform initialization | 0.9392±0.0011 | 0.9381±0.0007 | 0.939±0.0011 |
| xavier normal initialization | **0.9411 ± 0.0012** | 0.9389±0.0009 | 0.9393±0.0013 |
| batch normalization | 0.9401±0.0008 | 0.9389±0.0017 | **0.9394 ± 0.0011** |

Table S7: Our CNN+MLP models' AUPRC using different initializers or using batch normalization.

| | GM12878 | IMR90 | K562 |
|---|---|---|---|
| default initialization | **0.3752 ± 0.0038** | **0.3730 ± 0.0034** | 0.3701±0.0046 |
| xavier uniform initialization | 0.3717±0.0026 | 0.3691±0.0034 | 0.3697±0.0054 |
| xavier normal initialization | 0.3711±0.0043 | 0.3698±0.0023 | 0.3708±0.0029 |
| batch normalization | 0.3729±0.0025 | 0.3712±0.0027 | **0.3734 ± 0.0049** |

As shown in the table, adopting a different initializer or adding batch normalization did not significantly change the performance for our CNN+MLP models. We thus adopted the initializer and no batch normalization while training all our models.

#### 5.4 When training and testing Hi-C data are from different cell lines

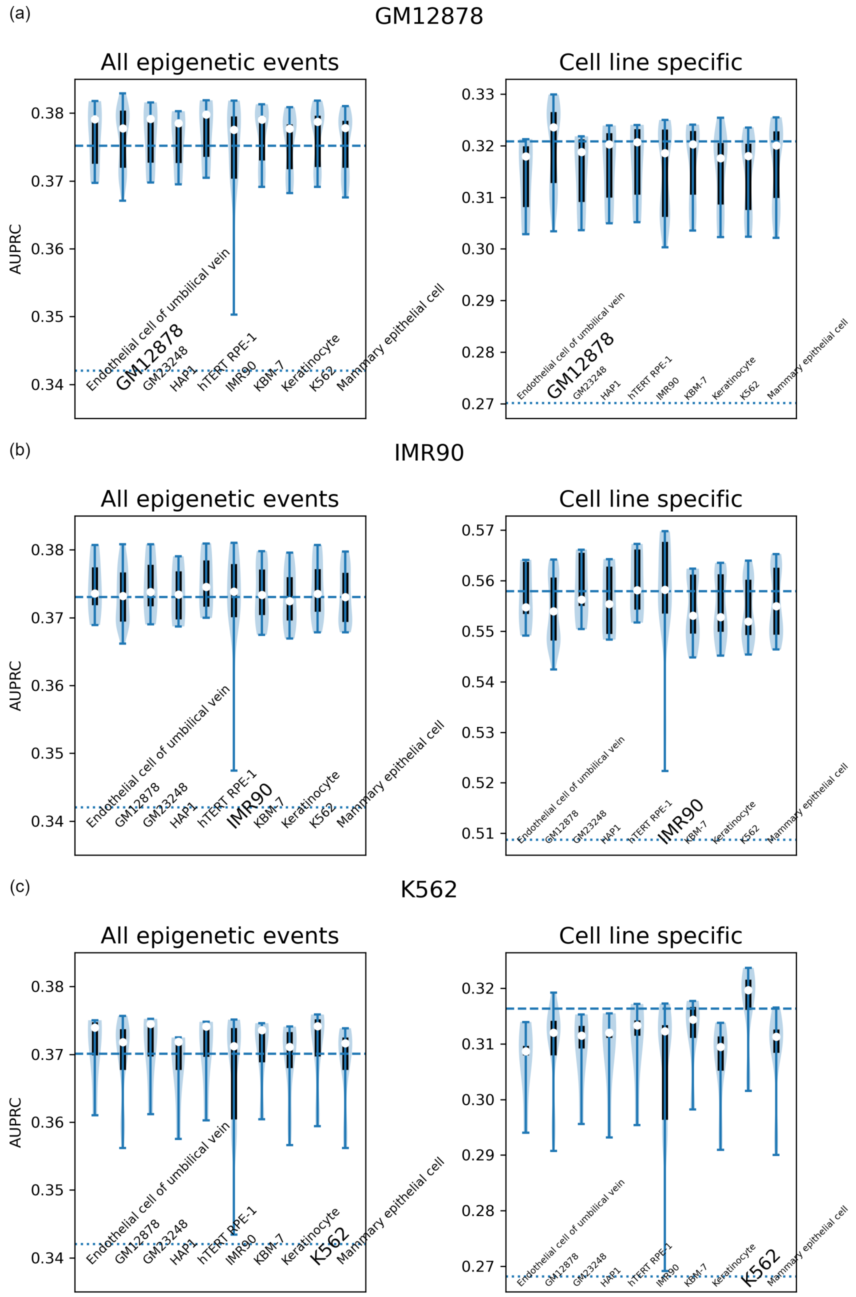


Figure S6: Violin plots of our CNN+MLP's performance when it is trained using Hi-C chromatin structure data for cell lines (a) GM12878, (b) IMR90, and (c) K562, respectively, but tested with other cell lines' Hi-C chromatin structure data (with corresponding name below each violin plot). Plotted are AUPRC distributions over five trials times (left) all epigenetic events or (right) those related to cell lines (a) GM12878, (b) IMR90, and (c) K562. The white dot, the box, and the whiskers indicate the median, the 25%–75% percentile, and the extremes, respectively. The dashed line represents the mean performance when each model is tested with the matched cell line's Hi-C data. The dotted line below represents the performance of sequence-only DeepSEA.

### 6 Extracting transcription factor (TF) binding motifs from learned epigenetic predictors

#### 6.1 Overall comparison across models

We report the numbers of convolutional kernels (out of 320, in the first layer of various models) that were found to match one or more known transcription factor binding motifs in Table S8. We also show the numbers of matched motifs in Venn diagrams in FigureS7. Note that our local DNA sequence embedding was based on CNN from DeepSEA or CNN/RNN from DanQ, where kernel sizes were 8 and 26, respectively. Thus directly comparing DeepSEA and DanQ (under the same E value cutoff in TOMTOM) would be unfair; and our comparisons are focused on either version without or with chromatin structure input (under various cell lines). Clearly with chromatin structure as input, our models showed complementarity to sequece-only models DeepSEA or DanQ.

Table S8: Number of kernels similar to known TF binding motifs.

| | GM12878 | IMR90 | K562 |
|---|---|---|---|
| DeepSEA (kernel size 8) | | 23 | |
| CNN (kerel size 8) + MLP | 25 | 24 | 25 |
| CNN+GCN (all 1 node feat.) | 27 | 28 | 25 |
| CNN+GCN (DNABERT node feat.) | 24 | 27 | 26 |
| DanQ (kernel size 26) | | 169 | |
| CNN (kernel size 26) /RNN + MLP | 167 | 164 | 166 |
| CNN/RNN+GCN (all 1 node feat.) | 167 | 166 | 166 |
| CNN/RNN+GCN (DNABERT node feat.) | 165 | 168 | 168 |

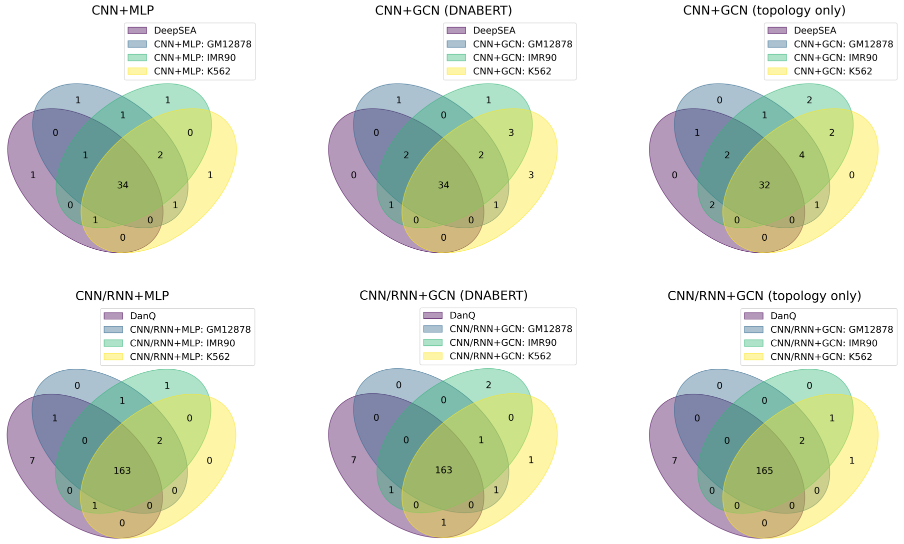


Figure S7: The numbers of known transcription factor binding motifs that were found a match to the sequence profiles in various models' learned convolutional kernels.

In the following subsections we analyze the specific motifs uniquely identified or missed by our models compared to DeepSEA or DanQ. We did not analyze our models when DNABERT was used because in such a case DNA sequence embedding occurs in both DNA sequence-embedding CNN and chromatin-embedding GCN and it would be unreasonable to only examine the first convolutional layer of CNN to extract motifs.

#### 6.2 Using CNN of kernel size 8 as in DeepSEA

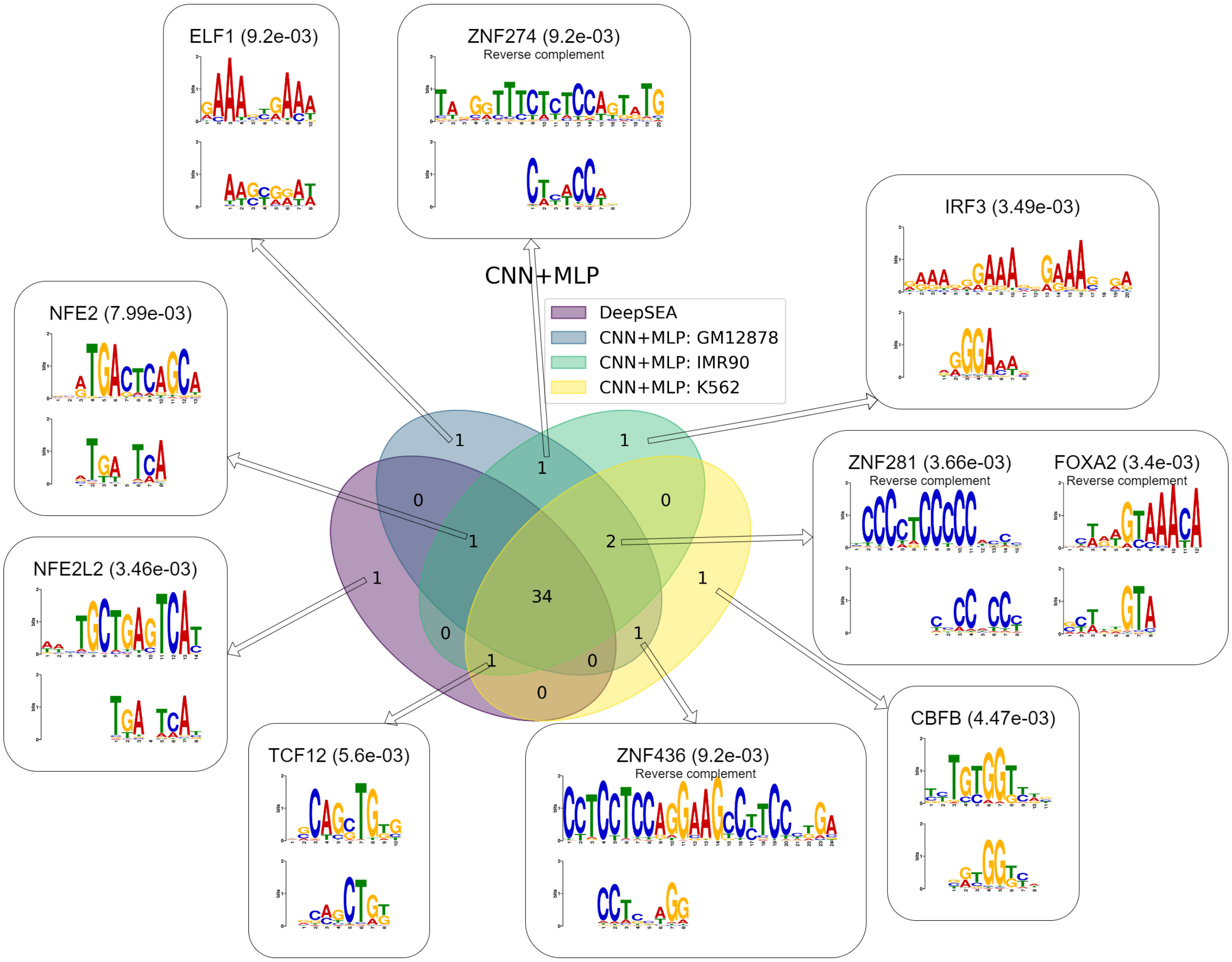


Figure S8: The numbers of motifs identified by CNN+MLP (using various cell lines' chromatin structures) versus DeepSEA.

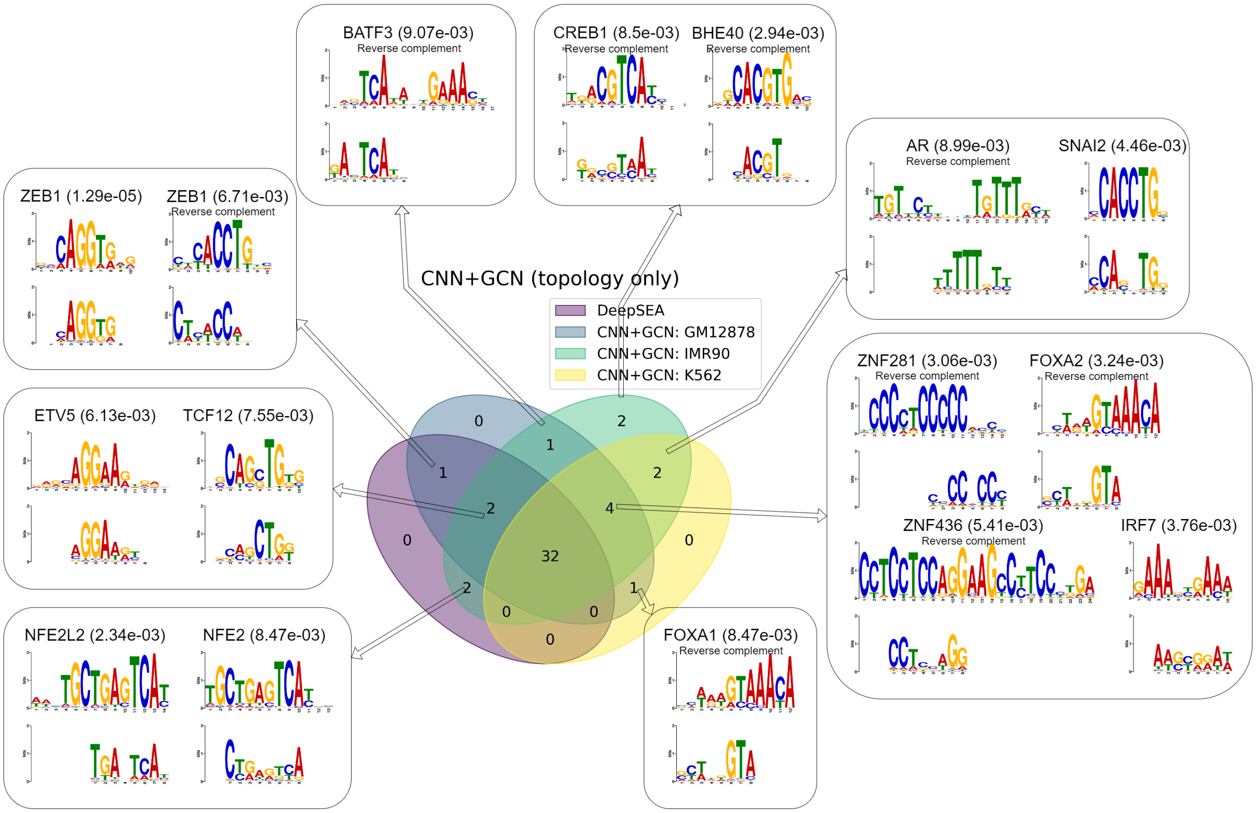


Figure S9: The numbers of motifs identified by CNN+GCN (topology only) (using various cell lines' chromatin structures) versus DeepSEA.

#### 6.3 Motif extraction using CNN of kernel size 26 as in DanQ

Our CNN/RNN+GCN consistently identified two motifs, TFs for genes POU3F1 (both strands) and TFDP1, that were missed by DanQ. The discovery of these two motifs was attributed to the use of chromatin structure that suggests long-range interactions. Specifically, POU3F1 (on Chromosome 1 or Chr. 1 in short) is a transcriptional activator that binds cooperatively with SOX4 (on Chromosome 6), SOX11 (Chr. 2) or SOX12 (Chr. 20). According to the chromatin topology defined by any of the 3 cell lines' interaction frequency matrix, POU3F1's region (100kbp resolution) is directly connected or separated by a common neighbor to SOX4, SOX11 and SOX12 regions. Without chromatin structures, the cross-chromosome TF cooperation would be much harder to identify from sequence alone. Similarly, TFDP1 (Chr. 13) encodes a transcription factor that functions cooperatively with E2F family members through the E2 recognition sites and plays a critical role in cell cycle regulation. According to the chromatin structure topology defined in the input Hi-C data, regardless of the cell line, the pseudo genes of TFDP1P1 (Chr. 1), TFDP1P2 (Chr. X), and TFDP1P3 (Chr. 15), as well as other genes important in cell cycle regulation including CCNA1 (Chr. 13), CCNA2 (Chr. 4), CCND1 (Chr. 11), CCND2 (on Chr. 12), CDK2 (Chr. 12), MYB (Chr. 6), E2F1 (Chr. 20), E2F2 (Chr. 1), E2F3 (Chr. 20) and CDC25A (Chr. 3) are all directly connected to or share a common neighbor with the TFDP1 region (100K-bp).

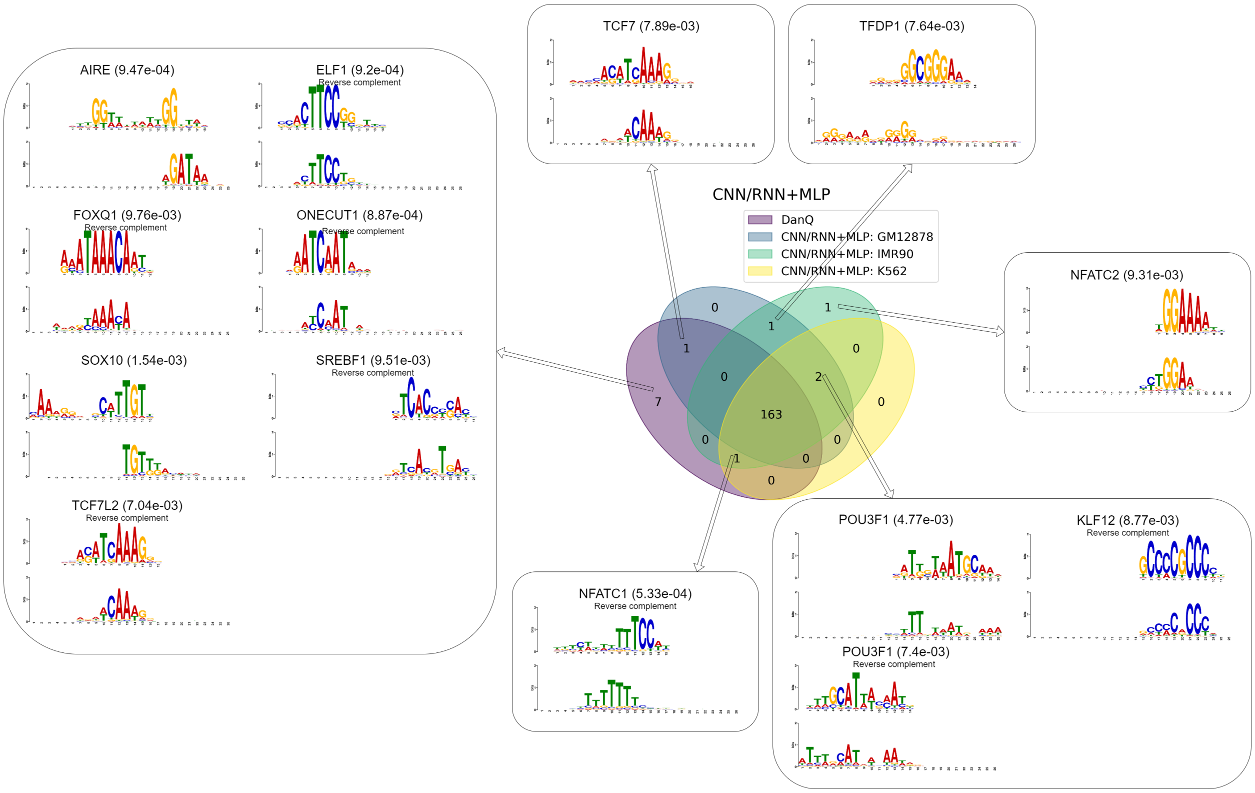


Figure S10: The numbers of motifs identified by CNN/RNN+MLP (using various cell lines' chromatin structures) versus DanQ.

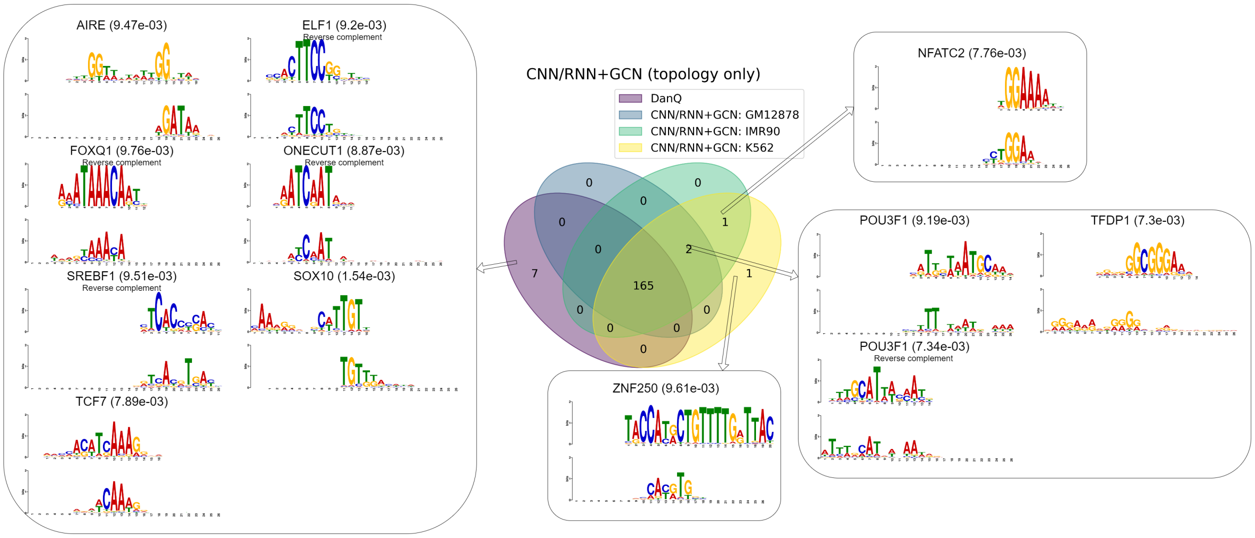


Figure S11: The numbers of motifs identified by CNN/RNN+GCN (topology only) (using various cell lines' chromatin structures) versus DanQ

#### 6.4 Understanding the false negatives

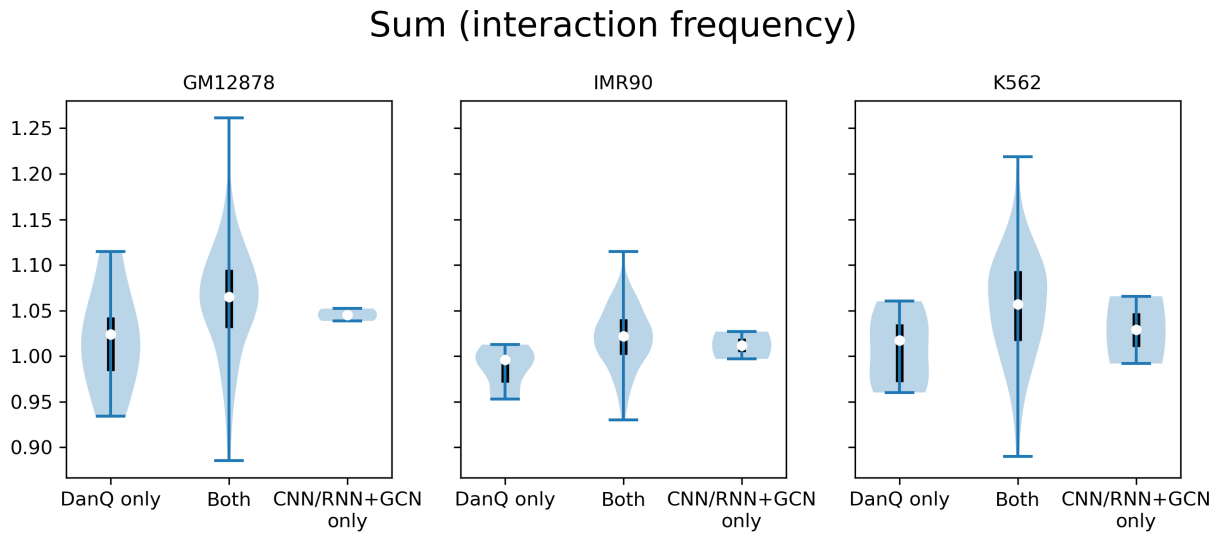


Figure S12: To have a better understanding of the motifs found by sequence-only DanQ but missed by our CNN/RNN+GCN topology only, we compared the sums of normalized interaction frequencies of the uniquely missed 7 genes, the commonly found genes, and the uniquely identified genes of our model.

### 7 Predicting noncoding variant effect: eQTL

We used the cell line-specific epigenetic predictions from our models to predict the eQTL's effect (expression level increasing or decreasing). Besides the performances of our basic CNN+MLP reported in the main text, we also report here the performances of CNN+GCN with DNABERT, CNN/RNN+MLP, and CNN/RNN+GCN with DNABERT.

L2 regularization term was tuned using validation loss and optimized as follows.

Regularization term used in 10-fold cross validation

| | DeepSEA | CNN+MLP | CNN+GCNw/DNABERT | DanQ | CNN/RNN+MLP | CNN/RNN+GCNw/DNABERT |
|---|---|---|---|---|---|---|
| GM12878 | 1E3 | 1E-3 | 1E1 | 1E-1 | 1E-1 | 1 |
| IMR90 | 1E-3 | 1E-3 | 1E-3 | 1E-3 | 1E-3 | 1E-3 |

As shown below, our chromatin structure-informed various models could outperform the corresponding sequence only model (DeepSEA or DanQ) in eQTL effect prediction, especially when the effects were strong.

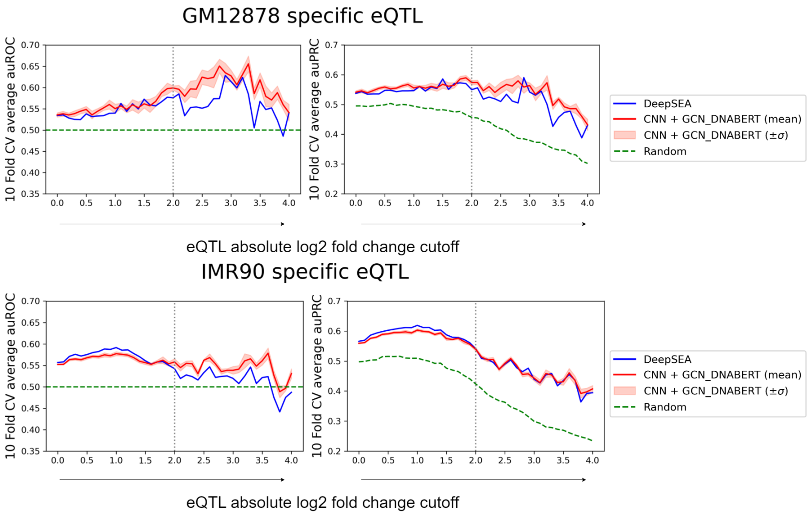


Figure S13: Using cell line specific epigenetic predictions from our CNN+GCN (with DNABERT to embed chromatin sequence and structure together) or DeepSEA to predict eQTL effect. For eQTLs with strong effect (the cutoff of expression log2 fold change at 2), our model for cell line GM12878 was 4.68$\sigma$ better (in AUROC) and 3.29$\sigma$ better (in AUPRC) than DeepSEA; and that for cell line IMR90 was 2.15$\sigma$ better (in AUROC) and 0.25$\sigma$ worse (in AUPRC) than DeepSEA. For the stronger effect eQTLs (expression log2 fold change cutoff at 2.5), our model for cell line GM12878 was 4.28$\sigma$ better (in AUROC) and 2.45$\sigma$ better (in AUPRC) than DeepSEA; and that for cell line IMR90 was 12.6$\sigma$ better (in AUROC) and 0.38$\sigma$ better (in AUPRC) than DeepSEA.

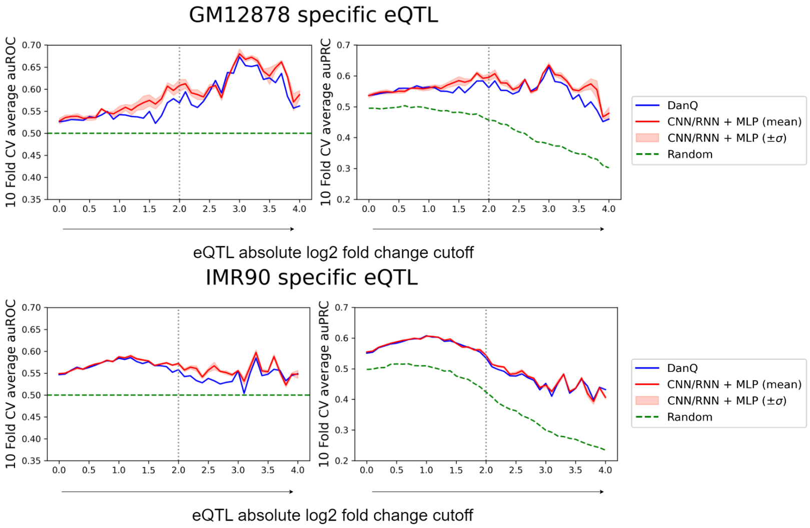


Figure S14: Using cell line specific epigenetic predictions from our CNN/RNN+MLP or DanQ to predict eQTL effect. For eQTLs with strong effect (the cutoff of expression log2 fold change at 2), our model for cell line GM12878 was 2.96$\sigma$ better (in AUROC) and 2.95$\sigma$ better (in AUPRC) than DanQ; and that for cell line IMR90 was 6.86$\sigma$ better (in AUROC) and 6.1$\sigma$ better (in AUPRC) than DanQ. For the stronger effect eQTLs (expression log2 fold change cutoff at 2.5), our model for cell line GM12878 was 1.39$\sigma$ better (in AUROC) and 1.36$\sigma$ better (in AUPRC) than DanQ; and that for cell line IMR90 was 3.89$\sigma$ better (in AUROC) and 1.38$\sigma$ better (in AUPRC) than DanQ.

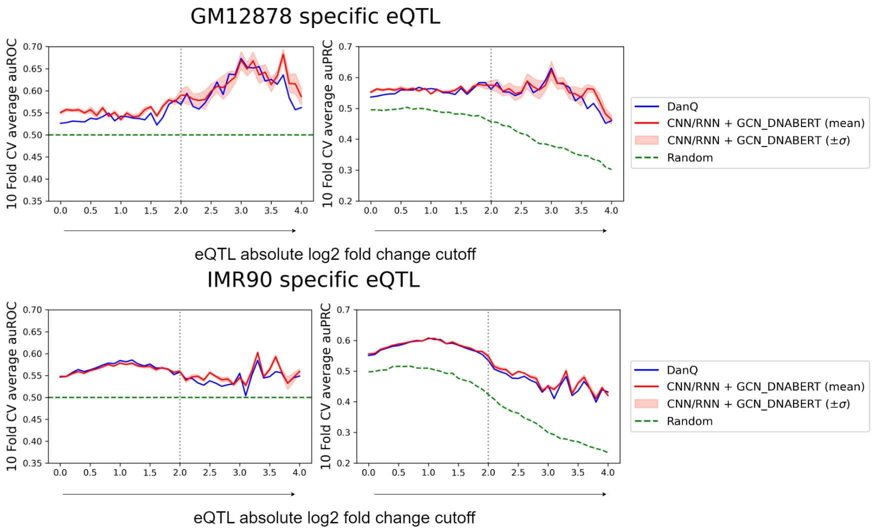


Figure S15: Using cell line specific epigenetic predictions from our CNN/RNN+GCN (with DNABERT to embed chromatin sequence and structure together) or DanQ to predict eQTL effect. For eQTLs with strong effect (the cutoff of expression log2 fold change at 2), our model for cell line GM12878 was 1.59$\sigma$ better (in AUROC) and 0.87$\sigma$ better (in AUPRC) than the DanQ; and that for cell line IMR90 was 0.56$\sigma$ better (in AUROC) and 2.09$\sigma$ better (in AUPRC) than DanQ. For the stronger effect eQTLs (expression log2 fold change cutoff at 2.5), our model for cell line GM12878 was 0.29$\sigma$ better (in auROC) and 0.18$\sigma$ better (in AUPRC) than DeepSEA; and that for cell line IMR90 was 8.47$\sigma$ better (in AUROC) and 8.78$\sigma$ better (in AUPRC) than DanQ.

### 8 Predicting noncoding variant effect: Pathogenicity

#### 8.1 Unsupervised "zero-shot" performances

We first report the unsupervised "zero-shot" learning performances on the test set curated from ncVarDB. No pathogenic/benigh variant data was used to re-train our models. Instead, absolute differences in epigenetic probabilities from pre-trained epigenetic predictors were used, in comparison to those from the sequence-only model DeepSEA.

Table S9: Unsupervised model prediction AUROC on ncVarDB: DeepSEA vs CNN+MLP

| | DeepSEA | Our GM12878 | Our IMR90 | Our K562 |
|---|---|---|---|---|
| Chromatin accessibility | 0.663 | 0.651 $\pm$ 0.003 | 0.688 $\pm$ 0.006 | 0.685 $\pm$ 0.002 |
| TF binding | 0.673 | 0.639 $\pm$ 0.007 | 0.668 $\pm$ 0.008 | 0.677 $\pm$ 0.005 |
| Histone mark modification | 0.683 | 0.700 $\pm$ 0.007 | 0.730 $\pm$ 0.004 | 0.728 $\pm$ 0.004 |
| Weighted across 3 types | 0.686 | 0.685 $\pm$ 0.006 | 0.718 $\pm$ 0.004 | 0.715 $\pm$ 0.003 |
| All features | 0.684 | 0.677 $\pm$ 0.006 | 0.709 $\pm$ 0.004 | 0.708 $\pm$ 0.002 |

Table S10: Unsupervised model prediction AUPRC on ncVarDB: DeepSEA vs CNN+MLP

| | DeepSEA | Our GM12878 | Our IMR90 | Our K562 |
|---|---|---|---|---|
| Chromatin accessibility | 0.155 | 0.151 ± 0.001 | 0.165 ± 0.002 | 0.164 ± 0.002 |
| TF binding | 0.173 | 0.167 ± 0.002 | 0.176 ± 0.003 | 0.182 ± 0.001 |
| Histone mark modification | 0.193 | 0.181 ± 0.005 | 0.210 ± 0.005 | 0.213 ± 0.009 |
| Weighted across 3 types | 0.173 | 0.169 ± 0.002 | 0.187 ± 0.00w | 0.189 ± 0.003 |
| All features | 0.176 | 0.173 ± 0.002 | 0.187 ± 0.002 | 0.190 ± 0.002 |

Table S11: Unsupervised model prediction AUROC on ncVarDB: DeepSEA vs CNN+GCN (DNABERT)

| | DeepSEA | Our GM12878 | Our IMR90 | Our K562 |
|---|---|---|---|---|
| Chromatin accessibility | 0.663 | 0.693 ± 0.003 | 0.695 ± 0.007 | 0.692 ± 0.005 |
| TF binding | 0.673 | 0.693 ± 0.005 | 0.677 ± 0.007 | 0.699 ± 0.001 |
| Histone mark modification | 0.688 | 0.701 ± 0.001 | 0.696 ± 0.005 | 0.704 ± 0.002 |
| Weighted across 3 types | 0.686 | 0.707 ± 0.003 | 0.703 ± 0.004 | 0.709 ± 0.002 |
| All features | 0.684 | 0.709 ± 0.004 | 0.701 ± 0.005 | 0.713 ± 0.002 |

Table S12: Unsupervised model prediction AUPRC on ncVarDB: DeepSEA vs CNN+GCN(DNABERT)

| | DeepSEA | Our GM12878 | Our IMR90 | Our K562 |
|---|---|---|---|---|
| Chromatin accessibility | 0.155 | 0.168 ± 0.003 | 0.172 ± 0.004 | 0.166 ± 0.003 |
| TF binding | 0.173 | 0.191 ± 0.002 | 0.177 ± 0.004 | 0.197 ± 0.002 |
| Histone mark modification | 0.193 | 0.182 ± 0.002 | 0.179 ± 0.005 | 0.176 ± 0.004 |
| Weighted across 3 types | 0.173 | 0.183 ± 0.002 | 0.184 ± 0.003 | 0.181 ± 0.002 |
| All features | 0.176 | 0.190 ± 0.002 | 0.186 ± 0.003 | 0.191 ± 0.002 |

Table S13: Unsupervised model prediction AUROC on ncVarDB: DanQ vs CNN/RNN+MLP

| | DanQ | Our GM12878 | Our IMR90 | Our K562 |
|---|---|---|---|---|
| Chromatin accessibility | 0.663 | 0.668 ± 0.002 | 0.687 ± 0.003 | 0.682 ± 0.002 |
| TF binding | 0.670 | 0.671 ± 0.005 | 0.689 ± 0.003 | 0.695 ± 0.002 |
| Histone mark modification | 0.671 | 0.719 ± 0.005 | 0.728 ± 0.002 | 0.727 ± 0.001 |
| Weighted across 3 types | 0.679 | 0.703 ± 0.003 | 0.718 ± 0.001 | 0.714 ± 0.002 |
| All features | 0.681 | 0.698 ± 0.004 | 0.713 ± 0.001 | 0.712 ± 0.001 |

Table S14: Unsupervised model prediction AUPRC on ncVarDB: DanQ vs CNN/RNN+MLP

| | DanQ | Our GM12878 | Our IMR90 | Our K562 |
|---|---|---|---|---|
| Chromatin accessibility | 0.152 | 0.153 ± 0.001 | 0.161 ± 0.002 | 0.157 ± 0.002 |
| TF binding | 0.164 | 0.172 ± 0.001 | 0.178 ± 0.002 | 0.184 ± 0.001 |
| Histone mark modification | 0.178 | 0.183 ± 0.004 | 0.195 ± 0.003 | 0.200 ± 0.004 |
| Weighted across 3 types | 0.167 | 0.173 ± 0.002 | 0.181 ± 0.001 | 0.181 ± 0.001 |
| All features | 0.168 | 0.176 ± 0.002 | 0.183 ± 0.001 | 0.185 ± 0.001 |

Table S15: Unsupervised models prediction AUROC on ncVarDB: DanQ vs CNN/RNN+GCN(DNABERT)

| | DanQ | Our GM12878 | Our IMR90 | Our K562 |
|---|---|---|---|---|
| Chromatin accessibility | 0.663 | 0.693 ± 0.002 | 0.687 ± 0.002 | 0.683 ± 0.002 |
| TF binding | 0.670 | 0.699 ± 0.003 | 0.685 ± 0.004 | 0.693 ± 0.002 |
| Histone mark modification | 0.671 | 0.688 ± 0.001 | 0.674 ± 0.002 | 0.681 ± 0.005 |
| Weighted across 3 types | 0.679 | 0.702 ± 0.002 | 0.691 ± 0.002 | 0.692 ± 0.003 |
| All features | 0.681 | 0.706 ± 0.002 | 0.693 ± 0.002 | 0.697 ± 0.002 |

Table S16: Unsupervised models prediction AUPRC on ncVarDB: DanQ vs CNN/RNN+GCN(DNABERT)

| | DanQ | Our GM12878 | Our IMR90 | Our K562 |
|---|---|---|---|---|
| Chromatin accessibility | 0.152 | 0.170 ± 0.002 | 0.168 ± 0.002 | 0.163 ± 0.001 |
| TF binding | 0.164 | 0.199 ± 0.002 | 0.181 ± 0.003 | 0.195 ± 0.001 |
| Histone mark modification | 0.178 | 0.168 ± 0.002 | 0.158 ± 0.002 | 0.163 ± 0.004 |
| Weighted across 3 types | 0.167 | 0.182 ± 0.002 | 0.176 ± 0.002 | 0.176 ± 0.002 |
| All features | 0.168 | 0.193 ± 0.003 | 0.182 ± 0.003 | 0.187 ± 0.001 |

#### 8.2 Supervised "few-shot" performances

In the main text, we showed that our Siamese neural network with pretrained epigenetic encoders, either CNN/RNN+MLP or CNN/RNN+GCN (with DNABERT), consistently further improve pathogenicity classification against zero-shot learning, using as few as tens of pathogenicity-labeled variants.

We additionally focused on the 535 variants from the ncVarDB test set that at least one of CADD, DANN, FATHMM-XF failed to make inference. These 535 variants are split based on their types: 305 substitution, 174 deletion, and 54 insertion variants. They are also split based on the genomic position: 358 intronic, 11 intergenic, 102 ncRNA, 26 3'UTR and 46 5'UTR variants. A summary is in Figure S16.

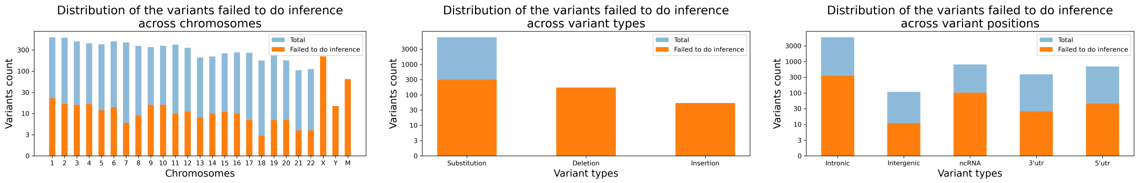


Figure S16: Summary of the 535 variants from ncVarDB that at least one of CADD, DANN, FATHMM-XF failed to make inference. Across all chromosomes, all variant types and variant positions, what proportion of the variants failed to get inference is shown in the figure.

Our models can make inference on the subset of test variants where SOTA models (such as CADD, DANN, or FATHMM-XF) failed to make predictions. So, our models are more generally applicable regarding the variant types and positions. Clearly, as shown in Figure 7 of the main text for all test variants, the performances for the subset of test variants in Figure S17 also had the upward trend while using more few-shot training examples. AUPRC values for this subset were better than those for the overall test set.

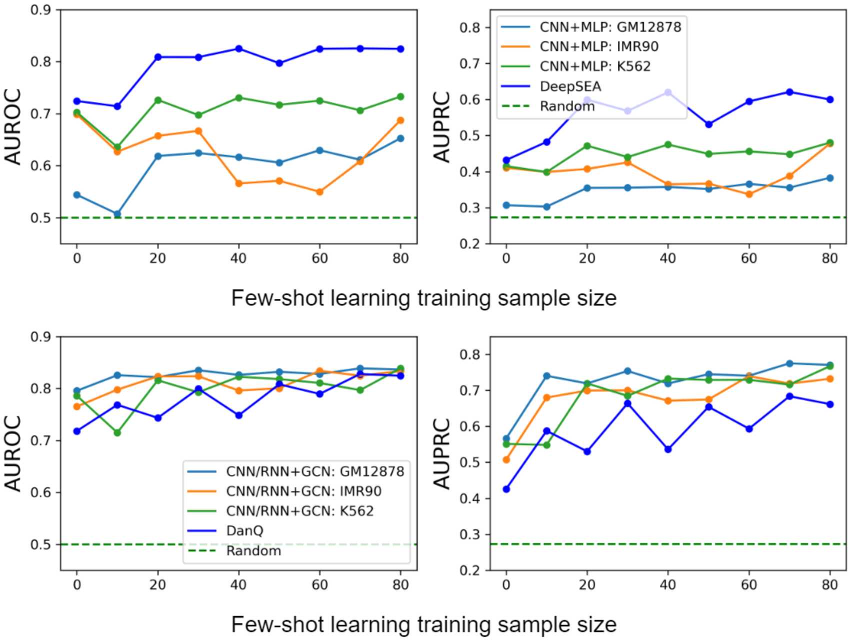


Figure S17: Our CNN+MLP (top) and CNN/RNN+GCN with DNABERT embedded node feature (bottom) prediction AUROC and AUPRC focusing on the subset that at least one of CADD, DANN, FATHMM-XF failed to make inference.

[5] Petar Veličković, Guillem Cucurull, Arantxa Casanova, Adriana Romero, Pietro Lio, and Yoshua Bengio. Graph attention networks. *arXiv preprint arXiv:1710.10903*, 2017.

[6] Yanrong Ji, Zhihan Zhou, Han Liu, and Ramana V Davuluri. Dnabert: pre-trained bidirectional encoder representations from transformers model for dna-language in genome. *Bioinformatics*, 37(15):2112–2120, 2021.

[7] Suhas SP Rao, Miriam H Huntley, Neva C Durand, Elena K Stamenova, Ivan D Bochkov, James T Robinson, Adrian L Sanborn, Ido Machol, Arina D Omer, Eric S Lander, et al. A 3d map of the human genome at kilobase resolution reveals principles of chromatin looping. *Cell*, 159(7):1665–1680, 2014.
